# SARS-CoV-2 reshapes m⁶A methylation in long non-coding RNAs of human lung cells

**DOI:** 10.1101/2025.06.08.658496

**Authors:** Cristina M. Peter, Caio O. Cyrino, Nilmar Moretti, Fernando Antoneli, Marcelo R. S. Briones

## Abstract

*N*⁶-Methyladenosine (m⁶A) is a key base modification that regulates RNA stability and translation during viral infection. While m⁶A methylation of host mRNAs has been studied in SARS-CoV-2-infected cells, its role in long non-coding RNAs (lncRNAs) is unknown. Here, we analyzed direct RNA sequencing (dRNA-seq) data from infected human lung cells (Calu-3) using a machine learning m⁶A detection framework. We observed a global increase in m⁶A levels across ten antiviral response– associated lncRNAs, with UCA1, GAS5, and NORAD—regulators of interferon (IFN) signaling— showing the most pronounced changes. This might, in part, explain the attenuated IFN expression observed in infected cells. We identified methylated DRACH motifs in predicted lncRNA duplex-forming regions, which may favor Hoogsteen base-pairing, which destabilize secondary structures and target interaction sites. These results provide new perspectives on how SARS-CoV-2 could impact lncRNAs to modulate host immunity and viral persistence through m⁶A-dependent mechanisms.

**In Brief:** Peter et al. show that SARS-CoV-2 infection alters m⁶A methylation of host lncRNAs by analysis of direct RNA sequencing data with machine learning. Key immune-regulatory lncRNAs show m⁶A alterations in RNA pairing regions, suggesting a novel mechanism by which m⁶A may modulate antiviral responses and promote viral persistence.

**Highlights:**

- Direct RNA-seq and machine learning reveal m⁶A changes in lncRNAs
- SARS-CoV-2 infection changes m⁶A patterns in antiviral response-associated lncRNAs
- LncRNAs UCA1, GAS5, and NORAD exhibit pronounced m⁶A remodeling upon infection
- m⁶A sites overlap RNA duplex-forming regions in immune-regulatory lncRNAs
- m⁶A may impact lncRNA duplexes via Hoogsteen pairing

**Graphical abstract:** 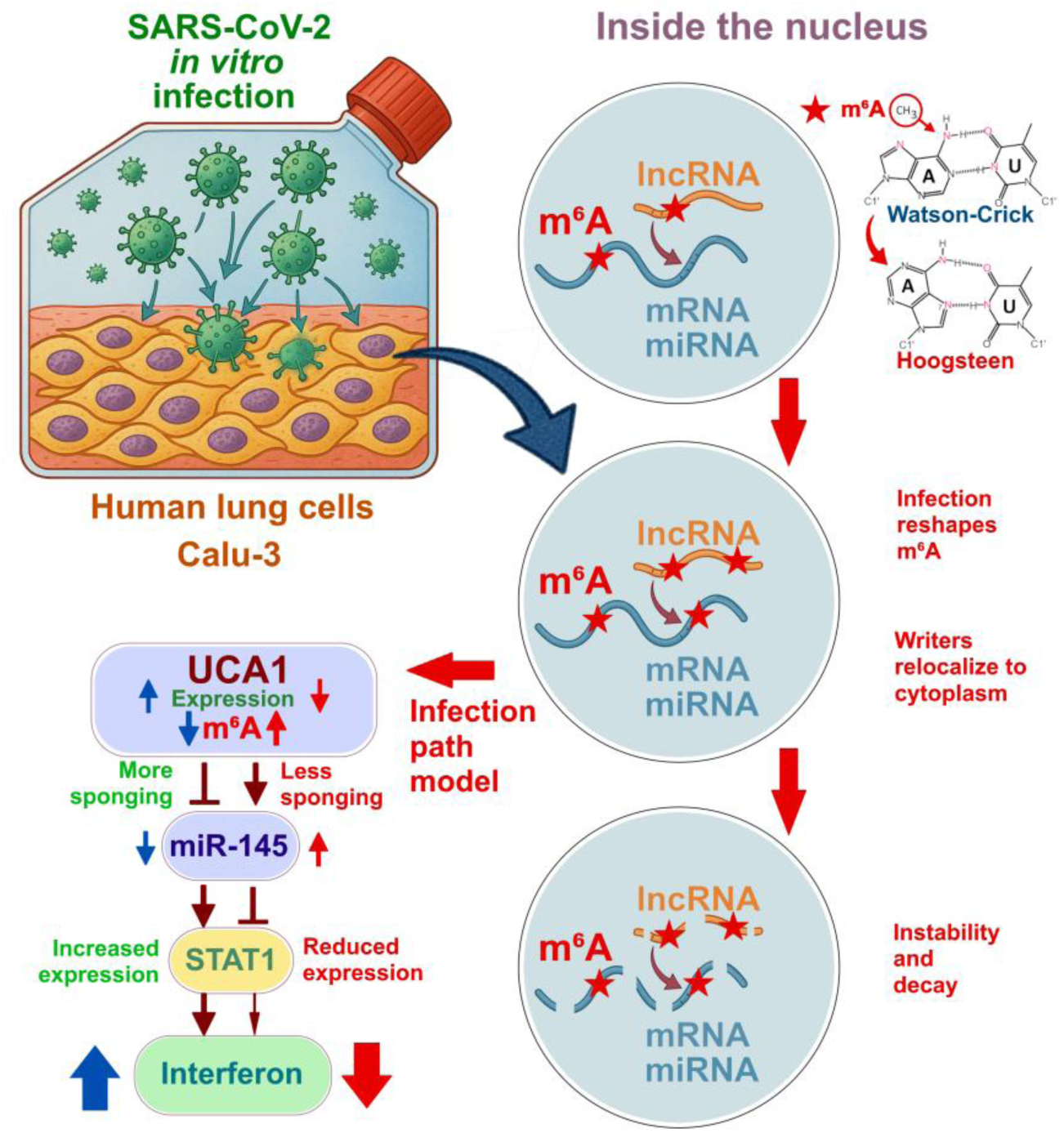

## INTRODUCTION

Severe acute respiratory syndrome coronavirus 2 (SARS-CoV-2), the etiological agent of COVID-19^1^, is a positive-sense single-stranded RNA virus that replicates in the cytoplasm of infected cells.^2,3^ In addition to viral RNAs, SARS-CoV-2 infected cells harbor diverse endogenous transcripts, including tRNAs, rRNAs, mRNAs, long non-coding RNAs (lncRNAs), and microRNAs (miRNAs), all of which may be affected by viral infection.^4,5^ SARS-CoV-2 disrupts host gene expression and post-transcriptional regulation,^6^ including the methylation landscape of host mRNAs via m⁶A, the most prevalent and functionally significant RNA modification in eukaryotic cells.^7^

Co-transcriptional m⁶A methylation occurs in the third position (adenosine) in DRACH motifs (D=A/G/U; R=A/G; A; C; H=A/C/U)^8^ by a methyltransferase complex composed of METTL3, METTL14, and WTAP (writers).^9^ This base modification is reversible, with FTO and ALKBH5 acting as demethylases (erasers),^10,11^ and is interpreted by m⁶A proteins such as YTHDFs and IGF2BPs (readers), which modulate RNA stability, splicing, translation, and decay.^12,13^ SARS-CoV-2 hijacks this machinery to enhance viral RNA stability and escape from immune detection.^5,14^ Notably, METTL3 interacts directly with the viral replication complex, facilitating m⁶A modification of viral RNAs and contributing to replication efficiency and IFN evasion.^15^

Long non-coding RNAs are at least 200 nucleotides long, do not contain coding regions and are emerging as key regulators of cellular homeostasis and immune responses.^16^ They function as molecular scaffolds, sponges, decoys, and guides for chromatin modifiers, RBPs, and miRNAs.^17–19^ In the context of SARS-CoV-2 infection,^20^ several lncRNAs are differentially expressed and implicated in cytokine modulation, IFN signaling, and viral genome processing.^6,21^ Yet, despite their importance, the epitranscriptomic regulation of lncRNAs during infection remains largely unexplored.

In particular, the extent to which m⁶A methylation shapes lncRNA function during SARS-CoV-2 infection is unknown. Given that lncRNAs often act through RNA-RNA or RNA-protein interactions, modifications that alter RNA structure could have significant regulatory consequences. Whether m⁶A methylation modulates such interactions—especially in immune-related lncRNAs—is a critical open question.^22^

Here, we examine m⁶A methylation dynamics in lncRNAs from SARS-CoV-2-infected Calu-3 cells, a human lung epithelial cell line that robustly supports infection and mirrors many features of *in vivo* airway epithelium.^23^ Using dRNA-seq data and a machine learning–based detection framework,^24^ we map high-confidence m⁶A sites across a curated set of immune-regulatory lncRNAs. We identify SARS-CoV-2–induced m⁶A remodeling in lncRNAs enriched at predicted RNA-RNA duplex regions and propose a novel hypothesis where m⁶A promotes transient Hoogsteen-like pairing^25^ in hairpins and base-pairing interfaces, potentially modulating lncRNA interactions and immune signaling.

## RESULTS

### Mapping dRNA-seq reads to lncRNAs

A curated set of 100 lncRNAs was assembled from published data based on their reported involvement in SARS-CoV-2 infection. Inclusion criteria encompassed associations with: (1) inflammation, (2) apoptosis, (3) biomarker potential, (4) innate immune response, and (5) adaptive immune response (**Table S1**). Direct RNA sequencing (dRNA-seq) reads were aligned to lncRNA reference sequences using Minimap2. From the initial set, 34 lncRNAs with higher read coverage were selected for downstream analyses, as greater read depth improves the reliability of methylation site detection (**Table S2**). BAM files corresponding to these 34 lncRNAs were analyzed for m⁶A modifications using a multiple instance learning framework (m6anet). Among them, ten lncRNAs exhibited strong evidence of m⁶A methylation—defined as prediction probabilities exceeding 60%— predominantly in infected cells. These ten lncRNAs were selected for detailed downstream analyses, as summarized in **Table 1**.

**Table 1.** Set of immune-related lncRNAs expressed in uninfected Calu-3 cells (U) and SARS-CoV-2 infected Calu-3 cells (I). These lncRNAs are a subset of curated a lncRNA set of 100 lncRNAs based on their relevance in immune response and inflammation regulation in SARS-CoV-2 infection. A total of 940,040 reads from uninfected cells and 1,055,956 reads from infected cells were analyzed.

| lncRNA | Gene ID | Description | Target | Regulation | Mapped Reads (U) | Mapped Reads (I) | Reference |
| --- | --- | --- | --- | --- | --- | --- | --- |
| BISPR | ENSG00000282851 | BST2 IFN stimulated positive regulator | BST2 (Tetherin) | Cis | 13,594 | 15,728 | <sup>20</sup> |
| GAS5 | ENSG00000234741 | Growth arrest specific 5 | TLR4, IL-6, TNF-α, SIRT1 miR-155 | Trans | 2,247 | 2,655 | <sup>47</sup> |
| LINC00278 | ENSG00000231535 | Long intergenic non-protein coding RNA 278 | IL-6 TNF-α | Trans | 9,734 | 11,510 | <sup>49</sup> |
| LINC00511 | ENSG00000227036 | Long intergenic non-protein coding RNA 511 | STAT3 NF-κB | Trans | 14,194 | 16,440 | <sup>81</sup> |
| MIR155HG | ENSG00000234883 | MIR155 host gene | NF-κB, IRF7 IFNAR1 | Trans | 1,348 | 1,804 | <sup>20</sup> |
| NORAD | ENSG00000260032 | Non-coding RNA activated by DNA damage | PUM1, NF-κB (indirect), p53 | Trans | 3,587 | 4,467 | <sup>45</sup> |
| THRIL | ENSG00000280634 | HNRNPL related immunoregulatory lncRNA | TNF-α, IL-6, IL-1β | Trans | 3,787 | 4,323 | <sup>46</sup> |
| UCA1 | ENSG00000214049 | Urothelial cancer associated1 | IL-6, TNF-α, NF-κB (via miRNAs) | Trans | 137 | 129 | <sup>51</sup> |
| CASC2 | ENSG00000177640 | Cancer susceptibility 2 | NF-κB, IL-6, TNF-α | Trans | 12,819 | 15,102 | <sup>50</sup> |
| LINC01619 | ENSG00000257242 | Long intergenic non-protein coding RNA 1619 | FOXO1 miR-27a-3p | Trans | 14,926 | 17,445 | <sup>49</sup> |

The dRNA-seq reads of infected and uninfected samples were mapped to the ten selected lncRNA reference sequences. A total of 940,040 dRNA-seq reads from uninfected cells (U) and 1,055,956 dRNA-seq reads from infected cells (I) were analyzed. The expression profiles of selected lncRNAs associated with the NF-κB, IFN-Stimulated Gene (ISG), *FOXO1*, STAT3 pathways in uninfected and infected cells are summarized in **Table 1**. Most of the lncRNAs were predicted to act via *trans*-regulatory mechanisms, targeting genes related to inflammation, immune signaling, and antiviral defense. BISPR was the only *cis*-acting lncRNA, located adjacent to the ISG *BST2* (Tetherin).

### Expression of lncRNAs in SARS-CoV-2 infected human lung cells

The expression profiles of the ten lncRNAs detailed in **Table 1** were estimated from dRNA-seq data by normalization of reads mapped to the lncRNA reference sequences (**Figure 1**). Most of the selected lncRNAs displayed modest upregulation upon infection, with fold changes ranging from approximately 1.03 to 1.06. Among upregulated lncRNAs, GAS5, LINC00511, MIR155HG, THRIL, and NORAD stood out due to their roles in inflammatory signaling. GAS5 and NORAD, both regulators of NF-κB, exhibited increased expression in infected samples, supporting their role in host immune defense. Notably, MIR155HG exhibited the highest increase in expression (FC≈ 1.19), consistent with its known role in inflammatory and antiviral responses. NORAD also showed a marked upregulation (FC≈ 1.11). In contrast, UCA1 was the only lncRNA to show a substantial decrease in expression, with a fold change below 0.90.

**Figure 1.**
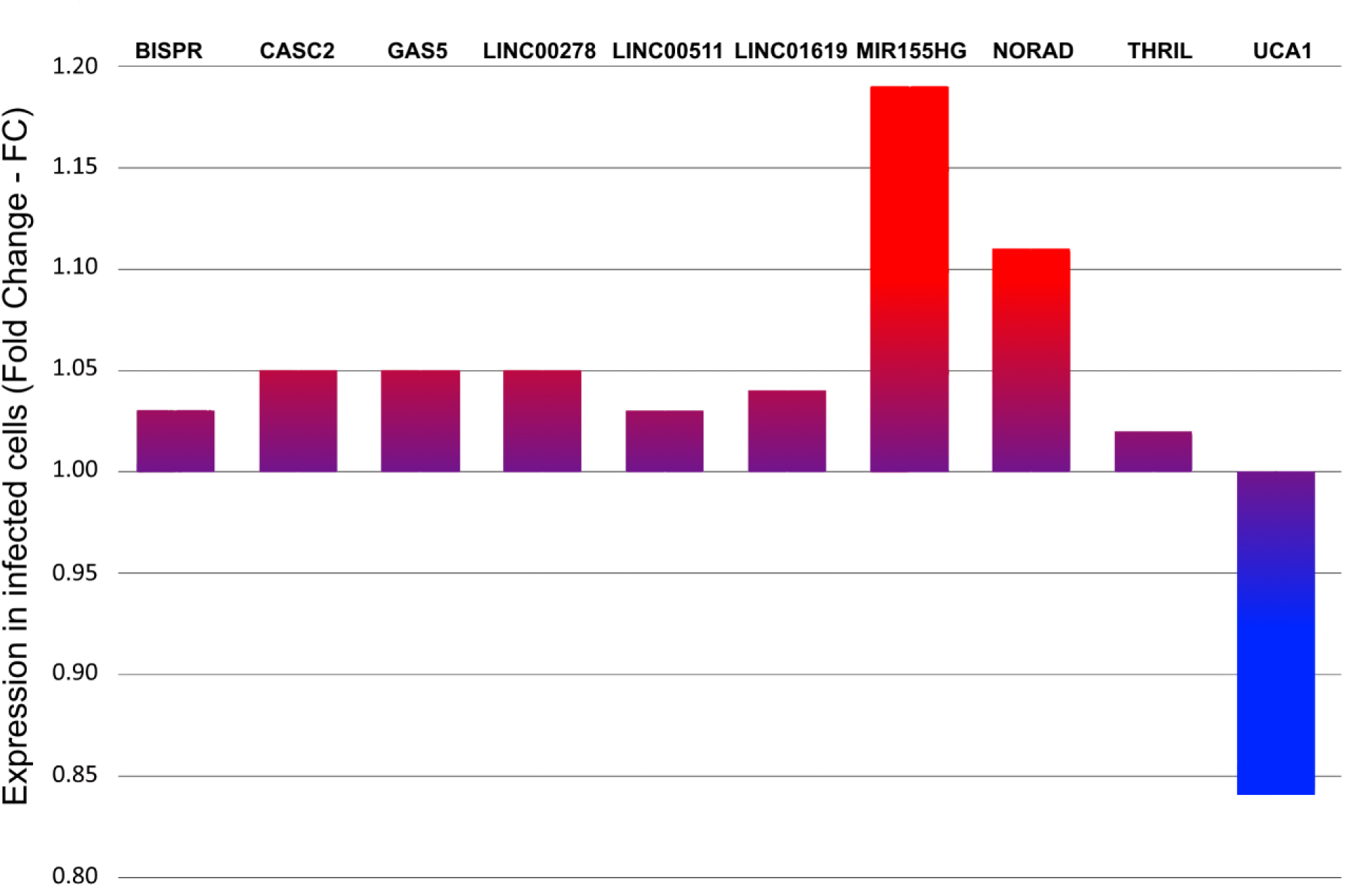
Expression of immune-associated lncRNAs in SARS-CoV-2-infected human cells (Calu-3). Bars represent the normalized fold-change (FC) of ten long non-coding RNAs (lncRNAs), associated with immune pathways, in SARS-CoV-2-infected cells as compared to uninfected cells. Values were derived from reference-based alignment of dRNA-seq data (BioProject PRJNA675370) and quantified using mapped read counts. Red indicates upregulation FC > 1.10, purple indicates 1.05>FC>1.00, and blue indicates downregulated (FC<1.00) in infected cells.

### Global changes of m⁶A patterns in SARS-CoV-2 infection

To assess whether the observed differences in m⁶A methylation between uninfected and infected samples were statistically significant, we performed a comparative analysis using the normalized methylation reads across all transcripts. This analysis shows the distribution of differentially methylated lncRNAs in Calu-3 cells (**Figure 2**). Each violin plot reflects the density of methylation values for uninfected and SARS-CoV-2-infected samples, with median values indicated by horizontal bars. The statistical significance of m⁶A differences between uninfected and SARS-CoV-2-infected Calu-3 cells, was assessed by the Wilcoxon–Mann–Whitney test (non-parametric) to compare normalized methylation signals across all detected sites in ten selected lncRNAs. The metric used, S= −log (standardized reads), represents the base-10 logarithm of the proportion of methylated sites normalized by read count.

**Figure 2.**
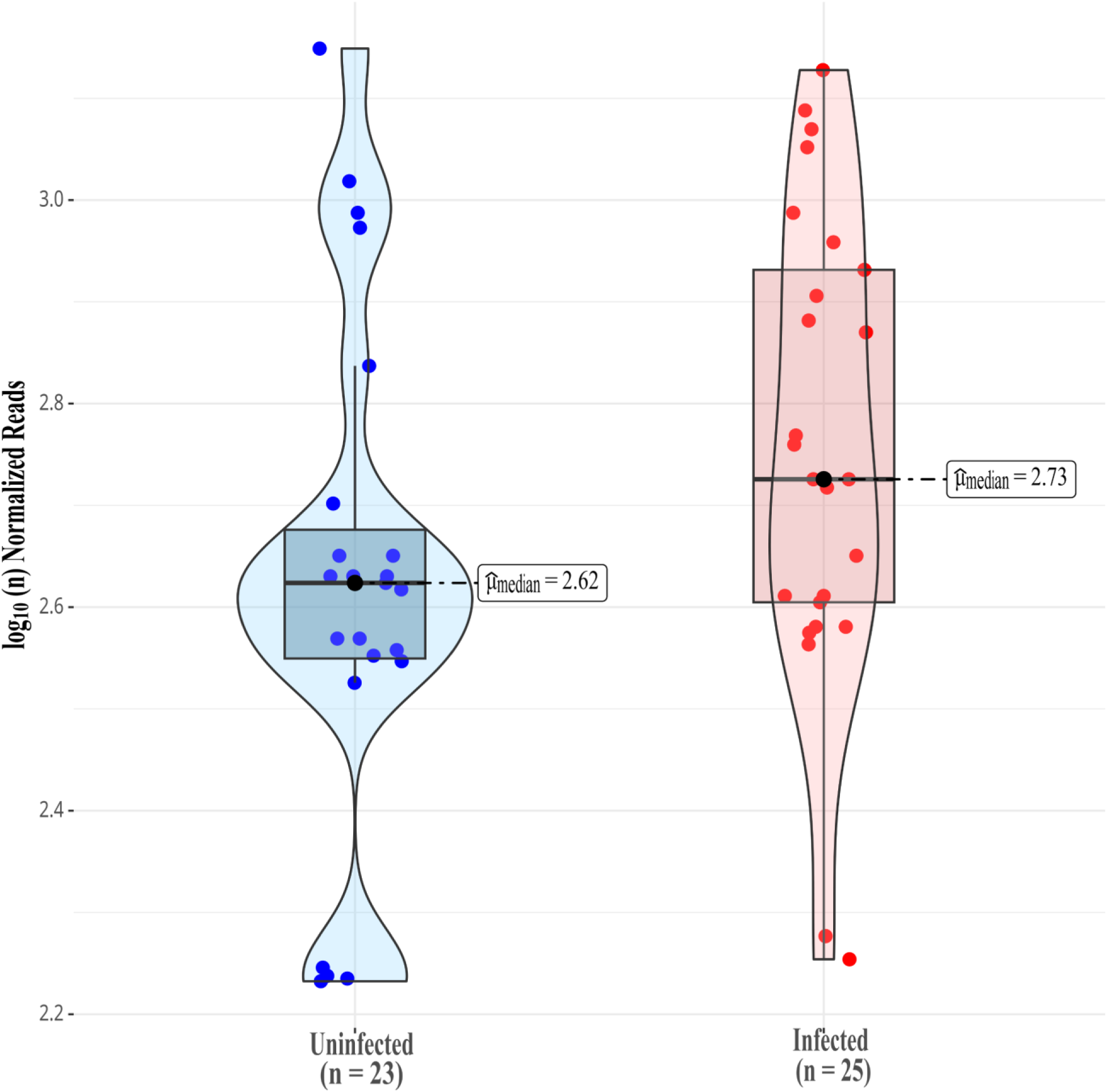
Global distribution of m⁶A sites in ten lncRNAs of SARS-CoV-2-infected and uninfected Calu-3 cells. The shaded areas of violin plots represent data distributions, and the horizontal bars indicate the medians. Methylation is estimated by multiple instance learning framework. Effect size and *p*-values were calculated using the Wilcoxon-Mann-Whitney test. The x-axis represents the sample groups, and the y-axis denotes S=−log (standardized reads), which is the base-10 logarithm of the proportion of methylated sites by the number of normalized reads in each dataset. The number of methylated sites is n=23 in uninfected cells and n=25 in infected cells with *W*_Mann-Whitney_=189.50, *p*=0.04, *ȓ*_biserial_=−0.34, CI_95%_=[−0.60, −0.02], *n*_obs_=48.

The violin plots reveal the distribution of m⁶A values, where each point represents an individual methylated site, and the shaded areas reflect the density of the data (**Figure 2**). The infected group (n=25) exhibited a slightly higher median methylation level (2.73) compared to the uninfected group (2.62), suggesting an overall increase in m⁶A modification upon infection. This difference is statistically significant (W= 189.5, *p*= 0.04), with a rank biserial effect size of −0.34, indicating a moderate shift in distribution. The 95% confidence interval (CI= [−0.60, −0.02]) further supports the reliability of this difference. The metric used, S= −log_10_ (standardized reads), captures the proportion of methylated sites relative to read-depth in each condition, allowing for normalization across samples.

### Mapping m⁶A modifications in lncRNAs

The regulatory potential of these lncRNAs, beyond their transcriptional changes, was assessed by their m⁶A epitranscriptome profiles. Using the same set of lncRNAs identified in expression analysis, we identified m⁶A modified sites with methylation probabilities ≥60% (**Table 2**). m⁶A sites were detected in the ten selected lncRNAs, including transcript coordinates, DRACH motifs, read counts, and methylation probabilities, revealing transcript-specific modification patterns (**Table 2**). Especially, several lncRNAs that exhibited transcriptional regulation, such as GAS5, THRIL, MIR155HG, and UCA1, also displayed consistent or increased m⁶A methylation signals following infection. The references used in these analyses were gene sequences so to include all exons of each lncRNA and avoid specificities of splicing variants expressed in different cell types and conditions.

**Table 2.** Distribution of m⁶A sites (*p*≥60%) in ten lncRNA genomic sequences in Calu-3 uninfected cells (U) and SARS-CoV-2 infected cells (I). m⁶A sites were identified using multiple instance learning as implemented in m6anet software. Not Detected (ND). Higher m⁶A methylation in infected cells (**) and uninfected cells (*). Bold fonts indicate lncRNAs with invariant m⁶A methylation. Numbering of m6A positions of the gene sequence refers to hg38 human genome assembly. Coverage is the number of reads that support the position. Probability is calculated by m6anet. K-mer is the specific DRACH motif where the m⁶A is located.

| lncRNA | m<sup>6</sup>A position | Coverage | Probability | K-mer |
| --- | --- | --- | --- | --- |
| BISPR (U) | 296* | 163 | 60% | UGACC |
| BISPR (I) | ND | ND | ND | ND |
| **CASC2 (U)** | **12758** | **164** | **78%** | **AGACU** |
| **CASC2 (I)** | **12758** | **157** | **79%** | **AGACU** |
| GAS5 (U) | 3550 | 20 | 83% | AGACA |
|  | 3979 | 63 | 82% | UGACA |
|  | 4062 | 76 | 80% | UGACC |
| GAS5 (I) | 3542** | 29 | 81% | GGACA |
|  | 3550 | 31 | 73% | AGACA |
|  | 3979 | 37 | 78% | UGACA |
|  | 4062 | 49 | 68% | UGACC |
| LINC00278 (U) | ND | ND | ND | ND |
| LINC00278 (I) | 11652** | 48 | 62% | GAACU |
| LINC00511 (U) | 3163* | 84 | 63% | AGACU |
| LINC00511 (I) | ND | ND | ND | ND |
| LINC01619 (U) | 7046 | 67 | 86% | AGACU |
|  | 7144* | 165 | 61% | AGACA |
| LINC01619 (I) | 7046 | 74 | 91% | AGACU |
| **MIR155HG (U)** | **9143** | **76** | **98%** | **AGACA** |
| **MIR155HG (I)** | **9143** | **54** | **96%** | **AGACA** |
| NORAD (U) | 2483* | 27 | 89% | AAACU |
|  | 2799 | 30 | 66% | GGACU |
| NORAD (I) | 2192** | 21 | 63% | AGACU |
|  | 2661** | 23 | 64% | GGACA |
|  | 2799 | 25 | 83% | GGACU |
| **THRIL (U)** | **615** | **160** | **95%** | **UGACU** |
| **THRIL (I)** | **615** | **149** | **95%** | **UGACU** |
| UCA1 (U) | 2367* | 29 | 91% | GGACC |
|  | 5738 | 56 | 91% | AGACA |
|  | 5922 | 66 | 93% | GGACC |
|  | 5971* | 79 | 70% | AGACC |
|  | 6098 | 68 | 77% | GGACA |
|  | 6158 | 66 | 77% | GGACA |
|  | 6197 | 66 | 91% | GGACA |
|  | 6285 | 80 | 61% | GAACU |
|  | 6293 | 78 | 90% | GAACU |
|  | 6390 | 63 | 76% | GGACA |
|  | 6683 | 41 | 86% | GAACU |
| UCA1 (I) | 5638** | 53 | 86% | GGACC |
|  | 5738 | 53 | 91% | AGACA |
|  | 5922 | 70 | 96% | GGACC |
|  | 6098 | 69 | 86% | GGACA |
|  | 6158 | 75 | 80% | GGACA |
|  | 6197 | 69 | 92% | GGACA |
|  | 6285 | 77 | 69% | GAACU |
|  | 6293 | 74 | 78% | GAACU |
|  | 6390 | 63 | 90% | GGACA |
|  | 6583** | 24 | 75% | AGACU |
|  | 6683 | 33 | 72% | GAACU |
|  | 6917** | 35 | 62% | UGACU |
|  | 6934** | 38 | 84% | UGACU |

Several lncRNAs exhibited distinct changes in their m⁶A methylation profiles in infected cells, including both the loss and gain of methylation sites (**Table 2**). BISPR, which showed upregulation at the transcriptional level, has a m⁶A methylated site (*p*= 60%) in uninfected cells, but no methylation sites were detected post-infection. Similarly, LINC00511 exhibited a methylated site with 63% probability in uninfected cells, whereas no detectable m⁶A signal was observed following infection. However, LINC00278, with no detectable methylation in uninfected cells, acquires a m⁶A-modified site (62%) upon infection.

Several lncRNAs exhibited invariant m⁶A modification patterns in uninfected vs infected cells comparison. THRIL displayed an invariant m⁶A site (with probability 95%) at the same position in both uninfected and infected cells, with a slight decrease in read coverage (**Table 2**). This suggests a robust and consistent epitranscriptome signature, potentially associated with maintenance of its role in TNFα regulation of inflammatory response. Similarly, CASC2 retained a methylation site at the same position in both conditions (78% and 79%) (**Table 2**). MIR155HG also maintained a highly methylated site (98% in uninfected cells and 96% in infected cells). Other lncRNAs showed m⁶A sites exclusively in infected cells, suggesting infection-induced epitranscriptome remodeling. NORAD exhibited two m⁶A modified sites in uninfected cells, and three in infected cells, including newly detected positions with methylation probabilities ≥60% (**Table 2**).

Several lncRNAs exhibited shared m⁶A-modified sites in both uninfected and infected cells, however with variations in methylation probability and/or read coverage, suggesting subtle post-transcriptional regulation rather than a complete gain or loss of modification. GAS5 retained multiple methylation sites in both conditions, although methylation probabilities decreased moderately in infected cells (e.g., from 83% to 73%, and from 82% to 78%), along with an overall decline in read counts (**Table 2**). NORAD, in addition to gaining new methylation sites, preserved at least one shared site (position 2799), with a methylation increase from 66% to 83% post-infection. Similarly, UCA1 displayed a mixed pattern, with numerous shared sites showing slight increases in methylation probabilities in infected cells (**Table 2**).

Individual methylation analysis of lncRNA transcripts of the most m⁶A-modified lncRNAs (**Table 2**) were performed with UCA1 (ENST00000397381.4) (**Table S3**), NORAD (ENST00000565493.1) (**Table S4**) and GAS5 (ENST00000430245.1) (**Table S5**). The transcript analysis confirms the higher methylation of UCA1 (2299 nt) with 48 DRACH motifs where 11 are methylated in infected cells and 12 in uninfected cells with probabilities ≥60% (*p* ≥60%) (**Table S3**). NORAD (5339 nt) also shows 48 DRACH motifs where 3 are methylated in infected cells and one in uninfected cells (*p* ≥60%) (**Table S4**). GAS5 (723 nt) has 12 DRACH motifs where only one is methylated with probability between 50%-60% in infected and uninfected cells (**Table S5**). The difference in GAS5 methylation when gene and transcript sequences are compared are due to methylated positions in introns which are identified in **Table 2** but that are not present in GAS5 transcript GAS5-027 that has predicted interaction regions with STAT2.

Given NEAT1 established role in viral infection response, we analyzed its main transcript (ENST00000501122.2, NEAT1-001, 22743 nt) (**Table S6**). NEAT1 is downregulated in dRNA-seq (FC= 0.72) and has only one m⁶A methylated position at 16,649 with probability 51.8% in infected cells and 77.3% in uninfected cells. All other DRACH motifs have methylation probabilities below 30%. NEAT1 is predicted to form RNA-RNA interactions with STAT1 and STAT2 (free energy ≤ –16 kcal/mol), which are key effectors in the interferon response pathway. The m⁶A methylated sites are not located in the target interaction regions.

### Changes in nucleotide preferences in DRACH motifs

We investigated whether the sequence context of m⁶A modified sites also varied in response to SARS-CoV-2 infection. Sequence logo analysis of methylated DRACH motifs shows alterations in nucleotide preferences in several positions (**Figure 3**). The analysis was performed by extracting each DRACH motif along with five flanking nucleotides on either side, providing a comprehensive view of the informational content and nucleotide bias surrounding methylated regions. Sequence profiles were generated separately for uninfected Calu-3 cells (**Figure 3A, 3C, 3E, 3G, 3I, 3K, 3M**) and SARS-CoV-2-infected cells (**Figure 3B, 3D, 3F, 3H, 3J, 3L, 3N**). It is important to note that only seven lncRNAs are represented, as three of the ten selected transcripts did not exhibit detectable m⁶A modifications in either condition. The y-axis indicates the information content in bits, where higher values reflect stronger sequence conservation and greater deviation from randomness.

**Figure 3.**
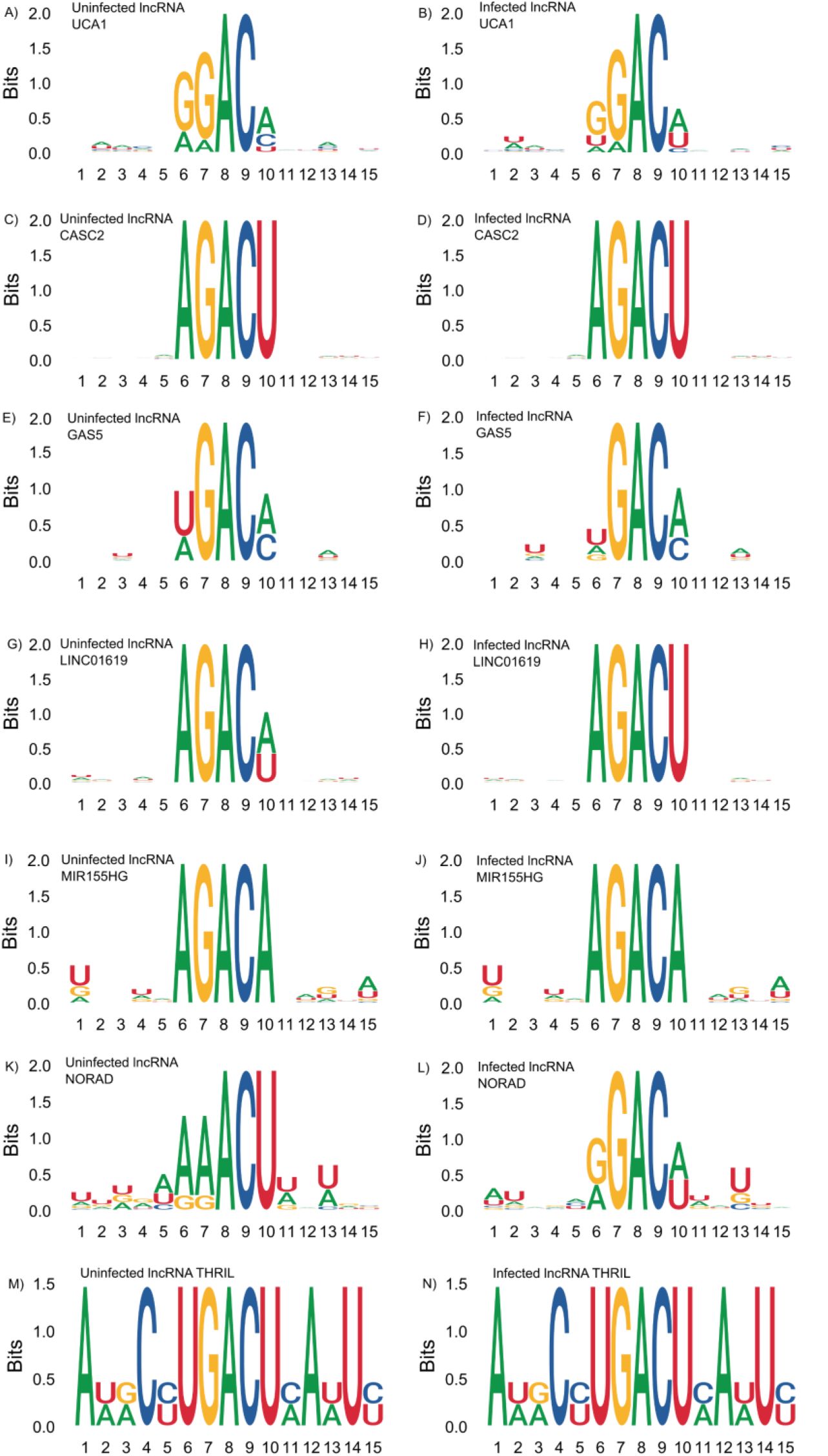
Nucleotide bias in lncRNA methylated DRACH motifs in SARS-CoV-2 infected cells. DRACH sequences containing predicted m⁶A sites (plus 5 nucleotides for each end) were aligned and stacked together to provide an overview of the informational content of methylated regions. Motif profiles in epitranscriptomes were obtained from Calu-3 Uninfected cells (A), and from samples of Infected Calu-3 cells (B). The ordinates indicate the score in Bits as it deviates from the null hypothesis (higher scores indicate stronger biases).

The core DRACH motif—particularly the GAC triplet—was highly conserved under both uninfected and infected conditions, although subtle variations emerged in flanking nucleotides, as summarized in **Table 3**. In several transcripts, such as GAS5 (**Figure 3E–F**), LINC00278 (**Figure 3G–H**), and LINC01619 (**Figure 3I–J**), we observed an increased frequency of adenine (A) and uracil (U) residues surrounding the DRACH motif in infected cells, suggesting a possible shift in sequence preference for m⁶A deposition during infection. Additionally, in lncRNAs like MIR155HG (**Figure 3K–L**) and CASC2 (**Figure 3C–D**), the information content (in bits) around the GAC core appeared slightly higher in infected cells, indicating stronger sequence conservation and potentially increased specificity of the m⁶A methylation machinery during SARS-CoV-2 infection. In contrast, BISPR (**Figure 3A–B**) maintained a highly similar sequence profile across both conditions, suggesting structural stability of its methylation motif. Finally, lncRNA THRIL revealed a conserved core DRACH motif (**Figure 3M–N**) in uninfected and infected cells, indicating canonical recognition by the methyltransferase complex. Notably, subtle variations in flanking nucleotide composition were observed upon infection, potentially reflecting context-specific adjustments in methylation targeting during viral stress.

**Table 3.**
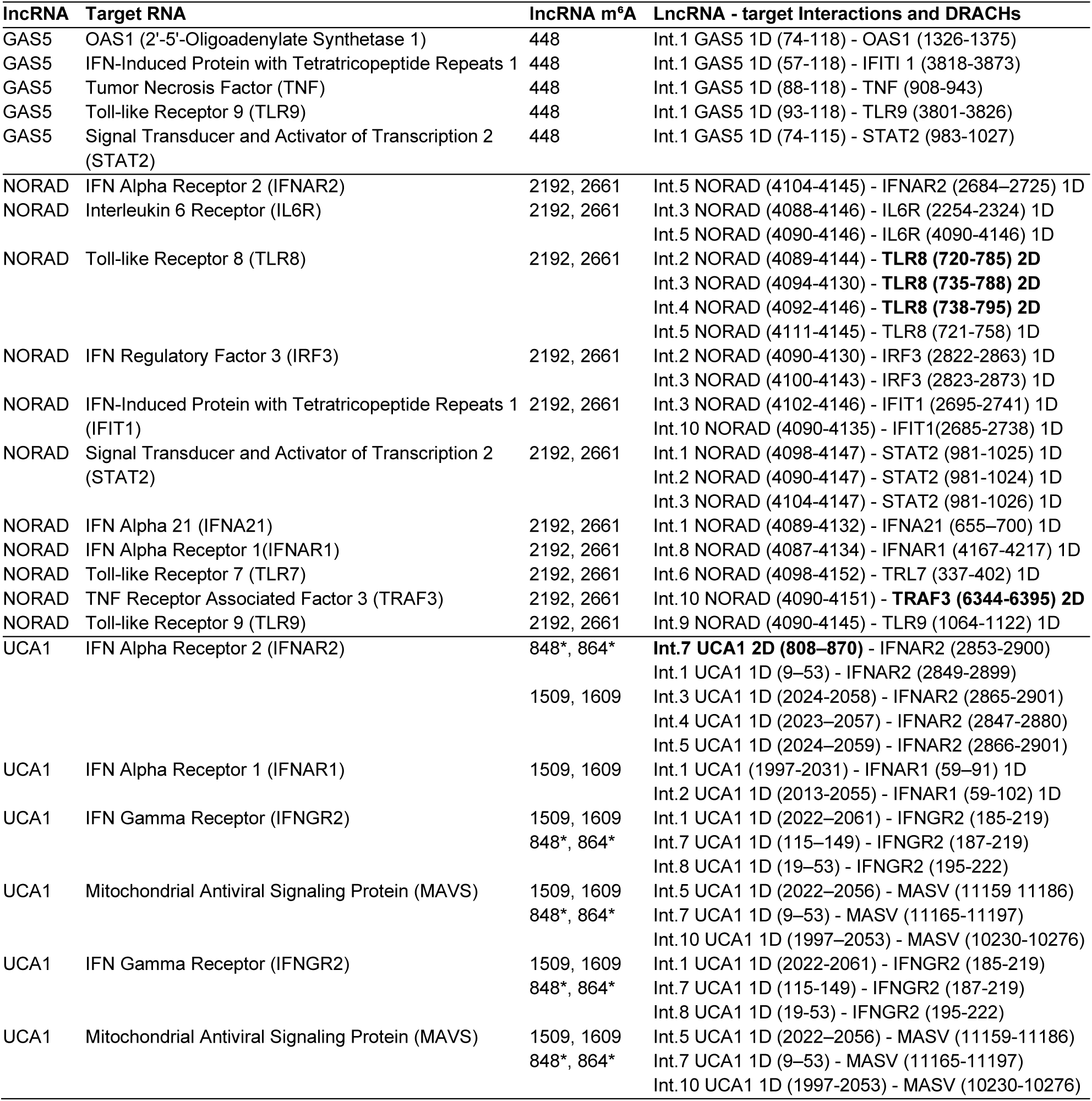
Positions of DRACH motifs, m⁶A sites and lncRNA-target RNA interactions in transcripts. Interactions (Int.) followed by the molecule (lncRNA or target) in which the DRACH was detected. Positions where m⁶A probability is within or in the vicinity of interaction region in transcripts GAS5 ENST00000430245 (GAS-027), NORAD ENST00000565493 (NORAD-001) and UCA1 ENST00000397381 (UCA1-001). Number of DRACH motifs in interaction regions, are indicated by, 1D=1 DRACH motif and 2D=2 DRACH motifs (bold font) and (*) indicate m⁶A methylated DRACH motifs within the interaction region, the other methylated sites are the closest to the interaction region.

**Table 4.** Summary table of the most and second-most frequent nucleotides at each position in the DRACH motif flanking m⁶A sites in each lncRNA under uninfected and infected conditions. The methylated base (A) is located at position 0. For each position, the dominant base is shown in uppercase; the second-most frequent base is shown in lowercase, separated by a slash. If two bases were equally frequent, both are shown in uppercase (e.g., A/U). Notable infection-associated shifts include the +2 position in UCA1 (A/c → A/u) and NORAD (U → A/U), suggesting infection-specific alterations in m⁶A sequence context.

| lncRNA | Condition | -2 | -1 | 0<br>(m<sup>6</sup>A) | +1 | +2 |
| --- | --- | --- | --- | --- | --- | --- |
| UCA1 | Uninfected | G/a | G | A | C | A/c |
| UCA1 | Infected | G/u | G | A | C | A/u |
| NORAD | Uninfected | A/g | A/g | A | C | U |
| NORAD | Infected | G/A | G | A | C | A/U |
| GAS5 | Uninfected | U/a | G | A | C | A/c |
| GAS5 | Infected | U/a | G | A | C | A/c |
| LINC01619 | Uninfected | A | G | A | C | A/U |
| LINC01619 | Infected | A | G | A | C | U |

### Positions of DRACH motifs and methylated m⁶A sites in lncRNAs

Methylated DRACH motifs were detected in the target recognition sites within UCA1 lncRNA (**Figure 4**). Also, differentially methylated DRACH motifs were found in lncRNAs secondary structure regions, highlighted in UCA1 (**Figure 5**). UCA1-001 (ENST00000397381) presented two methylated m⁶A sites (positions 848 and 864) within the predicted region of interaction with its target (positions UCA1-001 at 808–870 and IFNAR2-001 at 2853-2900) (**Table 3**, **Figure 5**). UCA1 shows m⁶A sites more methylated in infected cells, located within hairpins in the secondary structure where the m⁶A is in A-U pairs (**Figure 5**). The A-U pairs in which the A is a m⁶A could form, as discussed below, non-canonical base pairs that destabilize the RNA duplexes.

**Figure 4.**
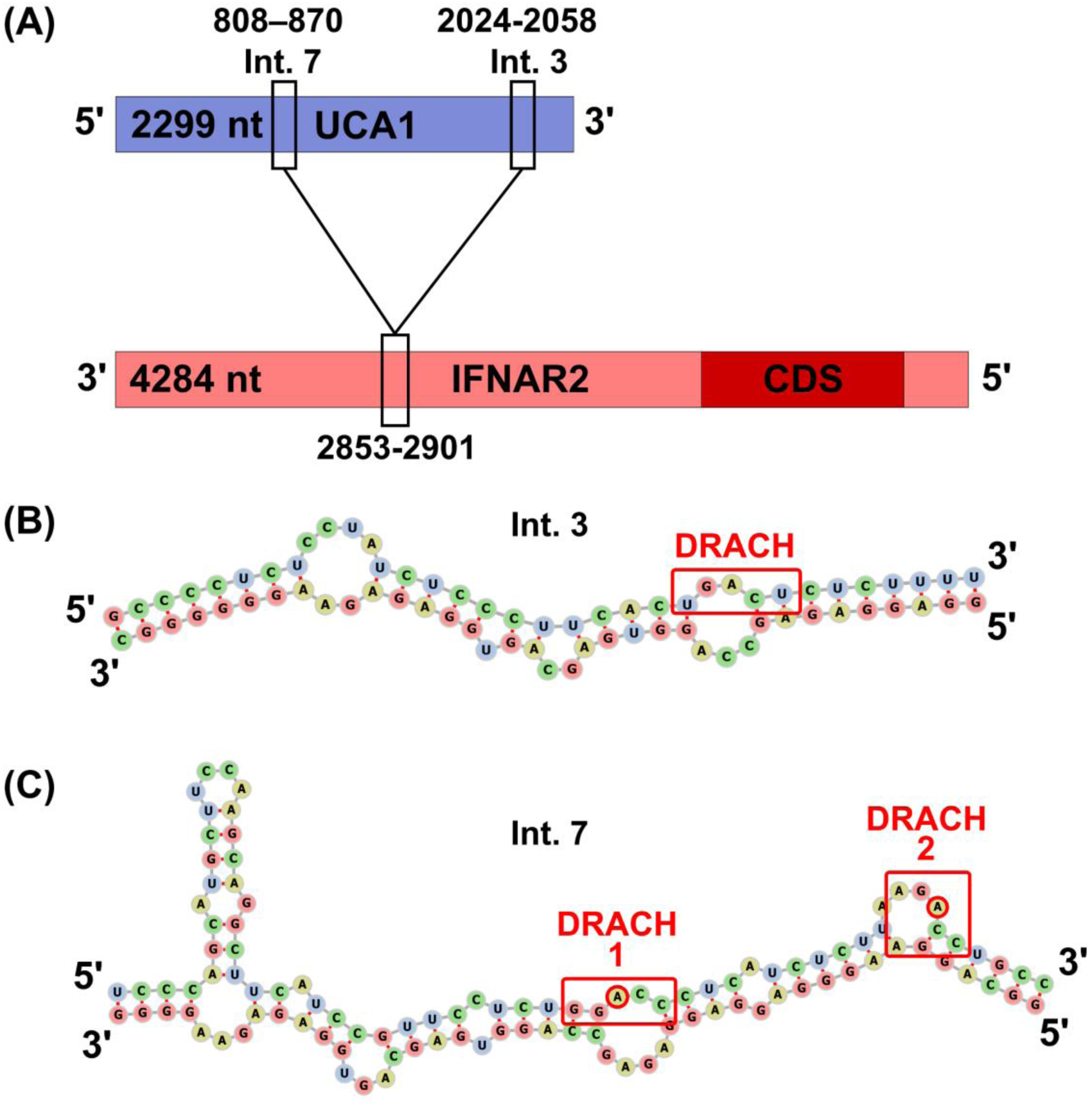
DRACH motifs in lncRNA UCA1 target interaction regions. (A) Map of the UCA1-IFNAR2 interaction regions 3 and 7. (B) The DRACH motif in interaction region 3 (Int. 3, free energy= −17.82 kcal/mol) with unmethylated A, also unpaired. (C) DRACH motifs 1 and 2 in interaction region 7 (Int. 7, free energy= −16.66 kcal/mol) contain methylated A bases, but they don’t pair with the target mRNA. UCA1-001 is the largest UCA1 splicing variant (ENST00000397381.4). IFNAR2 is the Interferon-alpha/beta receptor beta chain mRNA splicing variant IFNAR2-001 (ENST00000404220.3).

**Figure 5.**
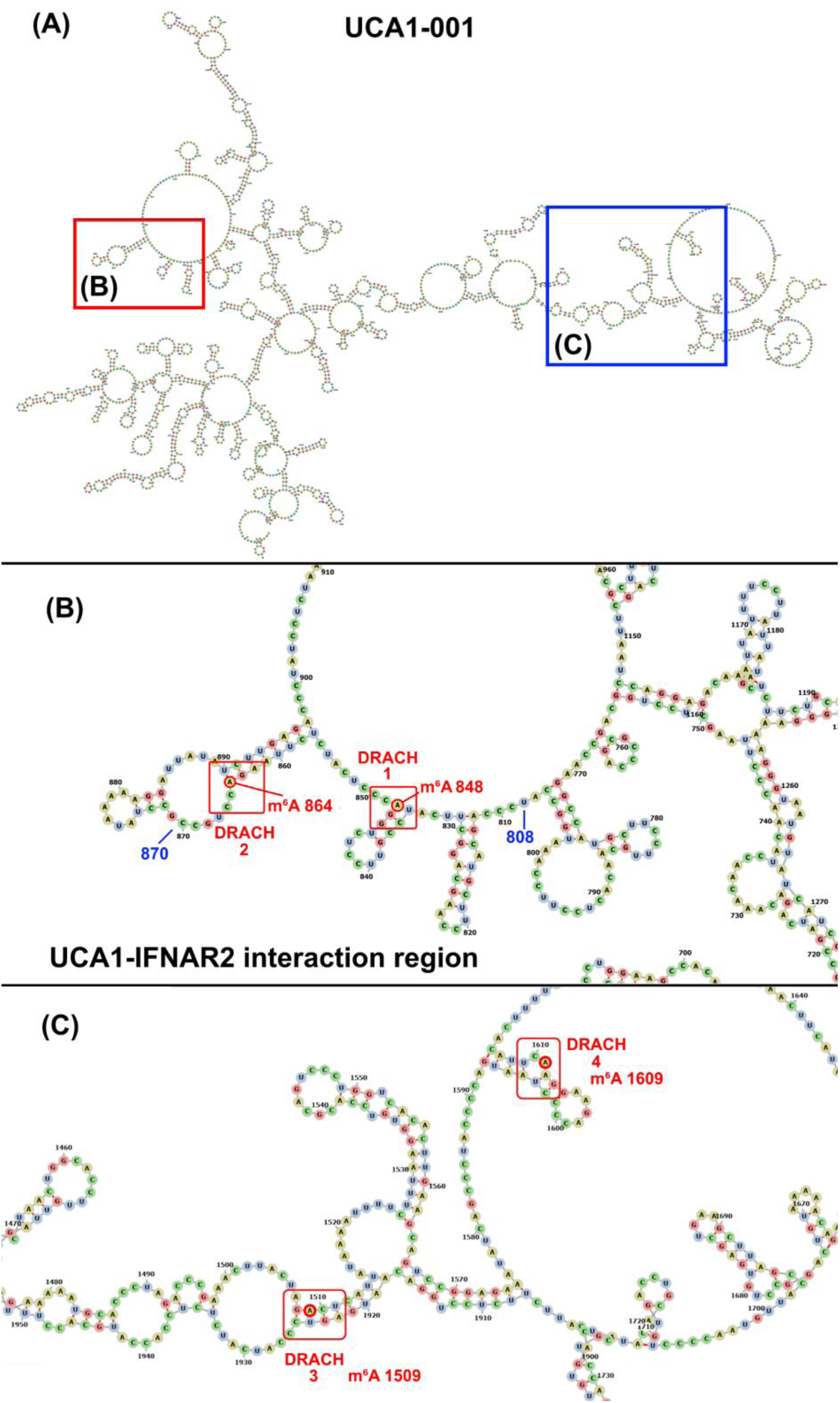
Location of DRACH motifs and m⁶A sites in the secondary structure of lncRNA UCA1. (A) Minimum free energy structure of the 2,229 nt transcript UCA1-001 (−563.20 kcal/mol) with hairpin loops detailed in (B) and (C). (B) Detail of the UCA1 region 808 to 870 that interacts with IFNAR2 mRNA. DRACH motifs 1 (GGACC) and 2 (AGACC) are in red boxes, m⁶A in 848 and 864 are indicated by red circles. (C) Detail of DRACH motifs with m⁶A modified 1509 that pairs with U (DRACH 3 - AGACU) and the unpaired m⁶A 1609 (DRACH 4 - GAACU).

In addition to target recognition regions and secondary structures, UCA1 is able to function as miRNA sponge. UCA1 transcript UCA1-001 (2299 nt) contains the 5’-AGCUGGAC-3’ motif that sponges miR-145 which in turn affects STAT1 expression and impacts IFN response. Another smaller sponging motif, 5’-CUGGAC-3’, is located in four positions in UCA1-001 transcript: (1) 508-513, (2) 844-849, (3) 1727-1732 and (4) 1912-1917. The sponging motif 844-849 overlaps with a m⁶A methylated DRACH motif (**Figure 6A**).

**Figure 6.**
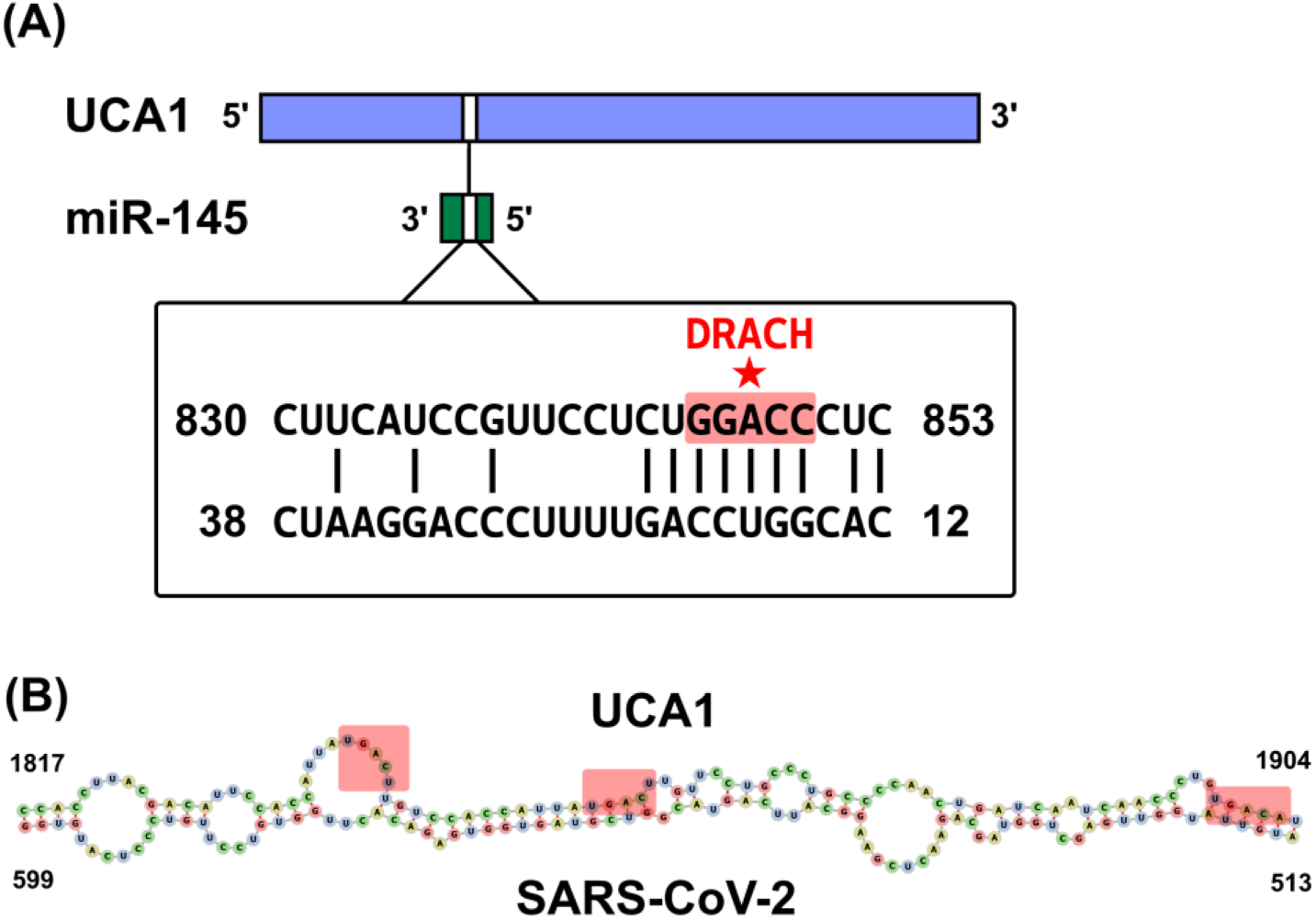
UCA1 interactions with miR-145 and SARS-CoV-2 RNA. (A) The UCA1 sponging motif has been experimentally demonstrated in its interaction with miR-145 in cancer and HCV infection contexts. The star indicates the methylated m⁶A site within the DRACH motif GGACC that overlaps with the sponging motif CUGGAC. The m⁶A pairs with U which could promote transient Hoogsteen pairs and potential duplex destabilization. (B) Interaction of UCA1 to SARS-CoV-2 genomic RNA. UCA1 positions 1817 to 1904 are predicted to interact with SARS-CoV-2 RNA at positions 513 to 599 with Energy −18.47 kcal/mol, Hybridization Energy −53.5 kcal/mol, Energy of SARS-CoV-2 = 27.17 kcal/mol and Unfolding Energy of UCA1 = 7.86 kcal/mol. Light red boxes indicate DRACH motifs. LncRNA-SARS-CoV-2 interactions were predicted using intaRNA program.

UCA1 is also capable at interacting with the SARS-CoV-2 genome RNA by forming a strong duplex in the region encompassing positions 1817 to 1904 that are predicted to interact with SARS-CoV-2 RNA positions 513 to 599 with Energy −18.47 kcal/mol, Hybridization Energy −53.5 kcal/mol, Energy of SARS-CoV-2 = 27.17 kcal/mol and Unfolding Energy of UCA1 = 7.86 kcal/mol as determined by intaRNA program (**Figure 6B**).

## DISCUSSION

### Global and local m⁶A changes in lncRNAs

The central idea of this study is to demonstrate that m⁶A methylation of lncRNAs is altered by SARS-CoV-2 infection. While global m⁶A remodeling of host mRNAs during SARS-CoV-2 infection has been previously reported, analogous studies of lncRNAs are lacking. Unlike coding transcripts, lncRNAs often act via structure-dependent interactions, making their methylation patterns—and the structural consequences thereof—particularly relevant.

We investigated the expression of ten selected lncRNAs, from an original curated set of 100 lncRNAs known to be associated with SARS-CoV-2 infection, particularly on the potential to modulate the host’s immune response, and viral replication. We observed a tendency of increased expression of these ten lncRNAs in SARS-CoV-2 infected Calu-3 cells, except for the downregulated UCA1, suggesting a coordinated activation of these transcripts as part of the host’s intracellular antiviral response (**Figure 1**). Also, the global pattern of increased lncRNA m⁶A methylation in infected cells, is consistent with increased m⁶A methylation in mRNAs, as previously observed^5^ (**Figure 2**). Although a global trend of increased lncRNA m⁶A methylation is statistically supported, given the lncRNA set considered (60% m⁶A probability cutoff), we observed that m⁶A methylation of each individual lncRNA and DRACH motif may increase, decrease or remain stable upon infection, depending on the context of each individual lncRNA and individual DRACH motif within each lncRNA, which we evaluated in detail (**Table 2**, **Table 3**, **Figure 2**).

### Expression levels and m⁶A patterns of lncRNAs and IFN response

According to changes in the expression levels of individual lncRNAs and their m⁶A methylation patterns we predict the outcome in terms of the IFN (INF) response as summarized in **Table 5**. Our analysis reveals distinct patterns of expression and m⁶A methylation among key long non-coding RNAs (lncRNAs) in SARS-CoV-2 infection (**Table 2, Tables S3, S4, S5 and S6**). Several lncRNAs with established or putative roles in IFN signaling exhibit coordinated changes in expression and m⁶A modification. UCA1 showed decreased expression but increased m⁶A methylation. Previous studies indicate that m⁶A on UCA1 reduces its ability to sponge miR-145-5p, thereby lifting repression on *SOCS7*, a known inhibitor of IFN signaling.^26^ This mechanism could result in suppressed IFN responses despite viral infection, aligning with known viral strategies to evade immune detection (**Table 5**). The simultaneous downregulation of UCA1 may reinforce this suppressive effect by reducing the lncRNA pool available for miRNA interaction. NORAD and GAS5 both exhibited increased expression and increased m⁶A levels. In NORAD, m⁶A has been shown to reduce sequestration of Pumilio RNA Binding Family Member (PUM1), a post-transcriptional repressor of IFN-stimulated genes (ISGs). This would enhance PUM1’s ability to degrade ISG transcripts, ultimately weakening the IFN response^27^ (**Table 5**). Similarly, in GAS5, m⁶A appears to reduce its sponging capacity for miRNAs like miR-21, which could diminish STAT1 activity and thereby attenuate IFN expression^28^ (**Table 5**). These patterns point to a broader trend in which m⁶A may facilitate viral immune evasion by functionally impairing lncRNA-mediated regulatory mechanisms, such as the IFN response.

**Table 5.** Prediction of IFN response as effect of lncRNAs expression (Exp.) and m⁶A methylation in SARS-CoV-2 infection.

| lncRNA | Exp. m<sup>6</sup>A |  | Proposed Mechanism | IFN | Ref. |
| --- | --- | --- | --- | --- | --- |
| UCA1 | ↓ | ↑ | m<sup>6</sup>A reduces miR-145-5p binding → ↑ SOCS7 → suppresses IFN signaling | ↓ | 26 |
| NORAD | ↑ | ↑ | m<sup>6</sup>A reduces PUM1 sequestration → ↑ PUM1-mediated decay of ISG mRNAs | ↓ | 27 |
| GAS5 | ↑ | ↑ | m<sup>6</sup>A reduces miRNA sponging (e.g., miR-21) → ↓ STAT1 expression | ↓ | 28 |
| BISPR | ↑ | ↓ | Reduced methylation enhances lncRNA stability/function → ↑ BST2 expression → activates IFN response | ↑ | 29 |
| CASC2 | ↑ | = | No change in m<sup>6</sup>A; IFN-modulatory role remains unclear | = | 30 |
| LINC00278 | ↑ | ↑ | m<sup>6</sup>A may affect translation of encoded micropeptides or RNA-RNA interactions; role in IFN unclear | Context-dependent | 33 |
| LINC00511 | ↑ | ↓ | Reduced methylation may enhance repressive function on IFN pathway | ↓ (if repressor) | 35 |
| LINC01619 | ↑ | ↑ | m<sup>6</sup>A likely affects structure or interactions; IFN pathway relevance not yet known | Unknown | 34 |
| MIR155HG | ↑ | = | Precursor of miR-155; no m<sup>6</sup>A change → likely unchanged miRNA processing or IFN effects | = | 31 |
| THRIL | ↑ | = | Known modulator of TNFα and IFNβ; unchanged m<sup>6</sup>A suggests conserved regulatory role | = | 32 |
| NEAT1 | ↓ | ↓ | ↓NEAT1 stability + ↓nuclear retention → ↓paraspeckles → less regulation of IRF7, RIG-I | ↓ | 82 |

In contrast, BISPR displayed increased expression but reduced methylation (**Table 2**). BISPR is a positive regulator of *BST2*, an antiviral factor. Hypomethylation may enhance BISPR stability or function, thus promoting *BST2* expression and activation of IFN responses^29^. This is one of the few examples in our dataset where a methylation change could support, rather than inhibit, antiviral immunity (**Table 5**). CASC2, while upregulated, did not exhibit a significant change in m⁶A methylation, and its role in the IFN pathway remains uncertain^30^ (**Table 5**). Similarly, MIR155HG and THRIL were upregulated without corresponding changes in methylation, suggesting that their regulatory roles—such as miR-155 biogenesis and modulation of TNFα/IFNβ expression—remain intact and are likely governed by transcriptional or post-transcriptional dynamics unrelated to m⁶A^31,32^ (**Table 5**). MIR155HG is a precursor of miR-155, a key regulator of innate immune response associated with hyperinflammatory conditions in COVID-19.^20^

LINC00278 and LINC01619 are both upregulated and hypermethylated but have ambiguous roles. LINC00278 encodes a micropeptide, and its m⁶A modification may influence translation or RNA–RNA interactions, though its impact on the IFN pathway appears context-dependent^33^ (**Table 5**). For LINC01619, structural or interaction-based modulation by m⁶A is plausible, but the relevance to antiviral immunity has yet to be clarified^34^ (**Table 5**). LINC00511 is upregulated but hypomethylated and may exert increased repressive effects on the IFN pathway if m⁶A typically constrains its function. This raises the possibility that demethylation could enhance its negative regulatory role under viral infection^35^ (**Table 5**).

NEAT1^36^ expression is elevated in saliva and nasopharyngeal samples of COVID-19 patients^37^ and upregulated in lung epithelial cells.^38^ The dRNA-seq here analyzed shows however, that NEAT1 is downregulated in infected Calu-3 cells (FC= 0.72) (**Table S2**). Mapping of dRNA-seq reads to NEAT1 identified DRACH motifs and a predicted m⁶A, above 60% probability, only in position 16648 with modest number or reads as compared to downstream sites (**Table S6**). This suggests that the NEAT1 transcript present in Calu-3 cells comprises exons located near the 3’ terminal region (approximately from positions 16000 to 18000) of the NEAT1 gene and not the ENSEMBL canonical 22k nt form (https://www.ensembl.org/). We also examined the location of this methylated site within predicted RNA secondary structures. These results suggest that NEAT1 may be subject to isoform-specific and structure-informed m⁶A regulation in SARS-CoV-2-infected cells Nevertheless, the combined effect of downregulated NEAT1 and low m⁶A methylation predicts an outcome of a weakened type I IFN response, potentially contributing to viral immune evasion (**Table 5**).

Our results provide a conceptual model based on lncRNA expression/methylation with testable predictions on IFN response. Accordingly, we examined the IFN expression profiles from the same dRNA-seq dataset, used for quantification lncRNA expression and m⁶A methylation, to test the anti-viral response predicted by our model (**Table 5**). We show that transcripts for IFN-α (IFNA1, IFNA2), and IFN-γ (IFNG) were undetectable in both conditions, echoing reports of suppressed or delayed type I and II IFN responses in SARS-CoV-2–infected cells^39^ (**Table 6**). Also, our data shows a selective induction of IFN-β (IFNB1), with 77 reads detected in infected cells versus none in uninfected cells, corresponding to a 68.55-fold change (**Table 6**). Although the absolute read count is modest, this finding is consistent with prior studies reporting low but inducible *IFNB1* expression in epithelial cells upon viral infection.^40^ Receptors for type I and II IFNs (IFNAR1, IFNAR2, IFNGR1, and IFNGR2) showed modest upregulation (fold changes ∼1.03–1.04), suggesting preserved or slightly enhanced signaling capacity. Together, these data align with a controlled, IFN-β–centered antiviral response and support our hypothesis that epitranscriptomic remodeling—including m⁶A modifications in lncRNAs—may fine-tune IFN signaling. Specifically, m⁶A enrichment in functionally relevant regions of lncRNAs such as UCA1, NORAD, and GAS5 may modulate their interactions with immune-related mRNAs, miRNAs, or RBPs, contributing to the specificity and magnitude of the host response.

**Table 6.** Expression of IFNs α, β and γ and respective receptors in uninfected and SARS-CoV-2 infected Calu-3 cells. The expression of IFNs and their receptors in infected cells (fold-change) inferred from dRNA-seq reads mapped to corresponding IFN and IFN receptor reference sequences.

| IFN/Receptor | Reads | Fold Change |
| --- | --- | --- |
| **IFN-α (*IFNA1*)** |  | **1** |
| Uninfected | 0 |  |
| Infected | 0 |  |
| **IFN-α (*IFNA2*)** |  | **1** |
| Uninfected | 0 |  |
| Infected | 0 |  |
| **IFN-β (*IFNB1*)** |  | **68.55** |
| Uninfected | 0 |  |
| Infected | 77 |  |
| **IFN-γ (*IFNG*)** |  | **1** |
| Uninfected | 0 |  |
| Infected | 0 |  |
| **IFNAR1 (IFN-α/$\beta$ receptor subunit 1)** |  | **1.03** |
| Uninfected | 14,996 |  |
| Infected | 17,393 |  |
| **IFNAR2 (IFN-α/$\beta$ receptor subunit 2)** |  | **1.04** |
| Uninfected | 12,646 |  |
| Infected | 14,815 |  |
| **IFNGR1 (IFN-γ receptor 1)** |  | **1.04** |
| Uninfected | 4,013 |  |
| Infected | 4,679 |  |
| **IFNGR2 (IFN-γ receptor 2)** |  | **1.03** |
| Uninfected | 16,603 |  |
| Infected | 19,201 |  |

**Table 7.**
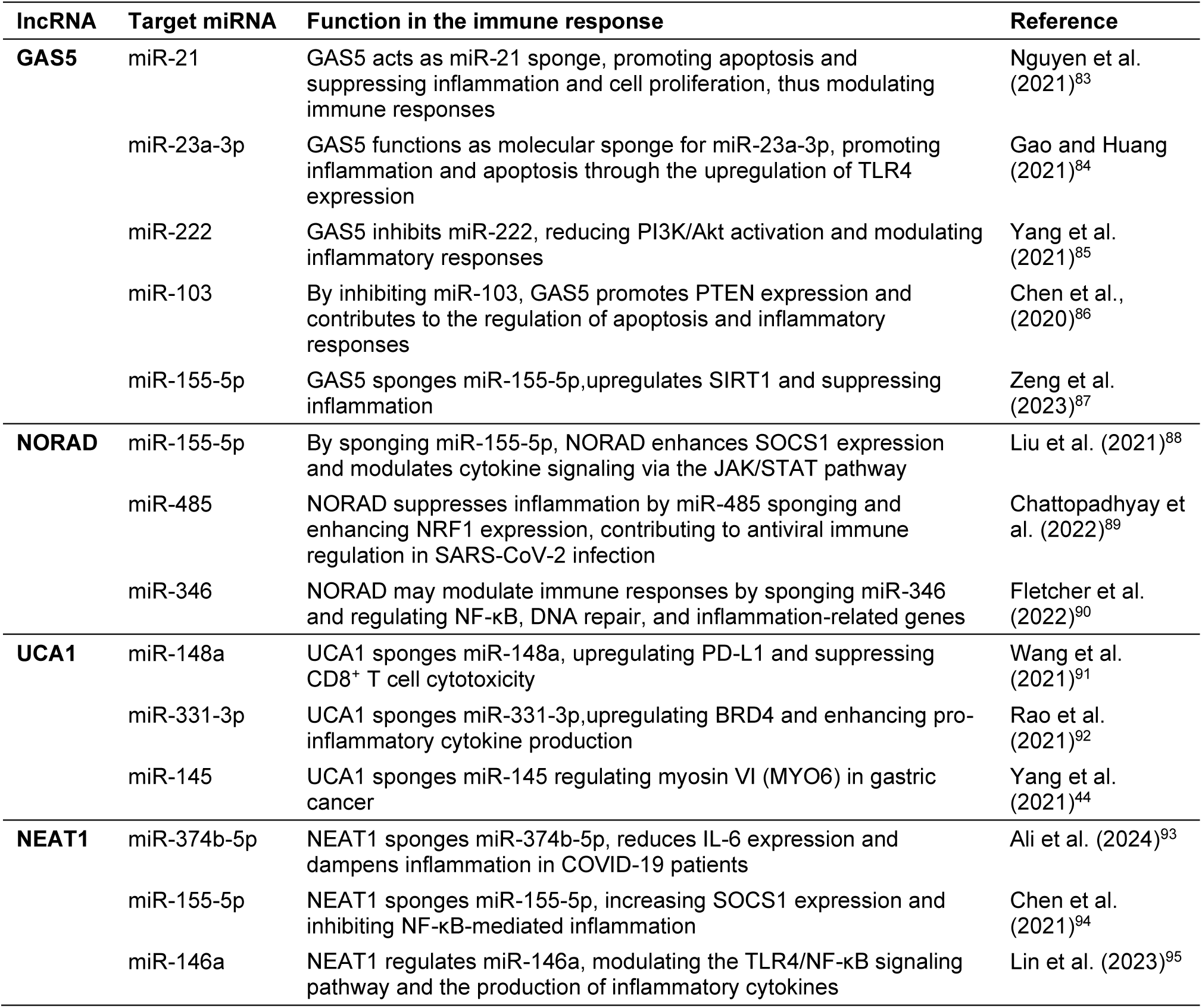
Immunoregulatory interactions of lncRNAs GAS5, NORAD, UCA1 and NEAT1 as sponges of key miRNAs.

| lncRNA | Target miRNA | Function in the immune response | Reference |
| --- | --- | --- | --- |
| **GAS5** | miR-21 | GAS5 acts as miR-21 sponge, promoting apoptosis and suppressing inflammation and cell proliferation, thus modulating immune responses | Nguyen et al. (2021)<sup>83</sup> |
|  | miR-23a-3p | GAS5 functions as molecular sponge for miR-23a-3p, promoting inflammation and apoptosis through the upregulation of TLR4 expression | Gao and Huang (2021)<sup>84</sup> |
|  | miR-222 | GAS5 inhibits miR-222, reducing PI3K/Akt activation and modulating inflammatory responses | Yang et al. (2021)<sup>85</sup> |
|  | miR-103 | By inhibiting miR-103, GAS5 promotes PTEN expression and contributes to the regulation of apoptosis and inflammatory responses | Chen et al., (2020)<sup>86</sup> |
|  | miR-155-5p | GAS5 sponges miR-155-5p, upregulates SIRT1 and suppressing inflammation | Zeng et al. (2023)<sup>87</sup> |
| **NORAD** | miR-155-5p | By sponging miR-155-5p, NORAD enhances SOCS1 expression and modulates cytokine signaling via the JAK/STAT pathway | Liu et al. (2021)<sup>88</sup> |
|  | miR-485 | NORAD suppresses inflammation by miR-485 sponging and enhancing NRF1 expression, contributing to antiviral immune regulation in SARS-CoV-2 infection | Chattopadhyay et al. (2022)<sup>89</sup> |
|  | miR-346 | NORAD may modulate immune responses by sponging miR-346 and regulating NF-κB, DNA repair, and inflammation-related genes | Fletcher et al. (2022)<sup>90</sup> |
| **UCA1** | miR-148a | UCA1 sponges miR-148a, upregulating PD-L1 and suppressing CD8<sup>+</sup> T cell cytotoxicity | Wang et al. (2021)<sup>91</sup> |
|  | miR-331-3p | UCA1 sponges miR-331-3p, upregulating BRD4 and enhancing pro-inflammatory cytokine production | Rao et al. (2021)<sup>92</sup> |
|  | miR-145 | UCA1 sponges miR-145 regulating myosin VI (MYO6) in gastric cancer | Yang et al. (2021)<sup>44</sup> |
| **NEAT1** | miR-374b-5p | NEAT1 sponges miR-374b-5p, reduces IL-6 expression and dampens inflammation in COVID-19 patients | Ali et al. (2024)<sup>93</sup> |
|  | miR-155-5p | NEAT1 sponges miR-155-5p, increasing SOCS1 expression and inhibiting NF-κB-mediated inflammation | Chen et al. (2021)<sup>94</sup> |
|  | miR-146a | NEAT1 regulates miR-146a, modulating the TLR4/NF-κB signaling pathway and the production of inflammatory cytokines | Lin et al. (2023)<sup>95</sup> |

### LncRNA m⁶A methylation and miRNAs

Several lncRNAs sponge miRNAs and lncRNA-miRNA interactions involve base pairing.^41^ For example, GAS5 sponges miR21 which in turn regulates the expression of STAT1, an activator of IFN transcription.^42^ The sequencing data here analyzed, however, does not contain any miRNAs because the default read size cutoff of Nanopore sequencing excludes reads smaller than 200nt. We attempted to detect miRNA in this dRNA-seq dataset using Minimap2, but no reads mapped to miRNA references with statistical significance. Therefore, we could not investigate whether miRNAs contain methylated DRACH motifs with the data at hand. This additional layer of gene regulation, involving the potential effects of m⁶A in miRNAs and the interaction to lncRNAs,^43^ remains an important question to be investigated.

Experimental evidence shows that sponging of miR-145 by UCA1 is dependent on sequence 5’-AGCUGGAC-3’ in UCA1 transcript EU334869.1 (ENST00000644174.2 UCA1-213 with 2688 nt) that forms a duplex with miR-145 sequence 3’-UUGACCUG-5’ in gastric cancer.^44^ This sponging motif, present in UCA1 transcript UCA1-213, is present in UCA1-001, here analyzed, at position 506-513. This motif does not contain a DRACH motif although it is close to the methylated m⁶A in DRACH in position 663. Therefore UCA1-001 with 2299 nt contains the 5’-AGCUGGAC-3’ motif and is predicted to sponge miR-145 which in turn affects STAT1 expression and impacts IFN response. How the differential m⁶A methylation of position 663 affects the sponging of miR-145 is to be tested experimentally. In HCV infection, UCA1 sponges miR-145-5p which regulates the level of the suppressor of cytokine signaling 7 (SOCS7), and in turn regulates the antiviral response in Huh7.5 cells.^26^ The UCA1 motif that mediates this sponging is 5’-CUGGAC-3’ that locates in four positions in UCA1-001 transcript; (1) 508-513, (2) 844-849, (3) 1727-1732 and (4) 1912-1917. The sponging motif 844-849 overlaps with a m⁶A methylated DRACH motif (**Figure 6A**). This intriguing observation offers an explicit hypothesis to be tested experimentally in future studies via genetic modification of Calu-3 cells with mutants of this sponge motif and their phenotypic changes in SARS-CoV-2infection.

Interestingly, the UCA1 region (1817–1904 nt) is predicted to strongly hybridize with the SARS-CoV-2 genomic RNA (513–599 nt; hybridization energy −53.5 kcal/mol) (**Figure 6B**). This region contains three DRACH motifs, although not methylated as indicated by in our analysis using m6anet. This unmethylated status may result from structural occlusion caused by early or persistent base-pairing with viral RNA, preventing access of the METTL3/METTL14 complex. Alternatively, the absence of m⁶A may facilitate hybridization, because methylation typically disrupts base pairing and promotes more open conformations. Both scenarios are consistent with the observed downregulation of UCA1 in infected cells and support a model in which the virus-induced RNA-RNA interaction alters the methylation landscape and post-transcriptional fate of host lncRNAs.

### LncRNAs and signaling pathways

NORAD^45^, THRIL^46^ and GAS5^47^ are involved in regulation of the NF-κB signaling pathway, a central axis of immune activation in response to respiratory viral infections and showed increased expression in infection (**Figure 1**). Although activation of the NF-κB pathway is essential for controlling viral infections, its excessive activation is associated with systemic inflammation and worse clinical outcomes in severe cases of COVID-19.^48^ Therefore, post-transcriptional and epigenetic regulation of lncRNAs such as GAS5 and NORAD may represent a compensatory mechanism of immune modulation, to balance antiviral defense and prevent tissue damage due to excessive inflammation. GAS5 acts as sponge, inhibiting miRNAs that repress components of the NF-κB pathway and amplify the inflammatory response.^47^ NORAD interacts with PUMILIO proteins (PUM1 and PUM2), key post-transcriptional repressors that bind to target mRNAs and promote their degradation and translational silencing. By sequestering PUMILIO proteins, NORAD prevents the repression of genes involved in DNA repair, stress responses, and cell cycle regulation.^45^ In the context of SARS-CoV-2 infection, this regulatory mechanism may contribute to genomic stability and modulate inflammatory responses. THRIL directly modulates TNF-α transcription by forming a complex with hnRNPL, a factor involved in inflammatory response.^46^ Therefore, the increased expression of these lncRNAs may play an active role in amplifying or modulating the cytokine-mediated inflammatory response during SARS-CoV-2 infection.

CASC2 and LINC01619 are downregulated in viral infections.^49,50^ However, these two lncRNAs show increased expression in SARS-CoV-2 infection as estimated from dRNA-seq data here analyzed (**Figure 1**). This suggests potential changes in RNA stability, alternative splicing, or other post-transcriptional mechanisms induced by viral stress. We also observed that most of the lncRNAs are *trans*-acting regulators, in line with their previously described roles in the regulation of multiple cellular pathways.^19^ An exception is BISPR, a *cis*-acting regulator of ISG *BST2* (Tetherin), whose coordinated co-expression was reported in COVID-19 patients,^20^ reinforcing its role in the IFN-mediated antiviral response. In viral infection, *trans*-acting lncRNAs can regulate processes ranging from ISG expression to redox homeostasis and apoptosis.^18^

Our analysis shows increased m⁶A methylation of GAS5, NORAD and UCA1 during infection, pointing to further layers of post-transcriptional regulation.^51^ The observed loss of the m⁶A modification in BISPR post-infection (**Table 2**) may indicate a decrease in m⁶A-mediated decay, therefore enhancing transcript stability and enabling its upregulation and cis-mediated regulation of *BST2*. In contrast, lncRNAs LINC00278 and NORAD acquired novel m⁶A sites post-infection, suggesting a potential role for m⁶A in stress-induced activation or stabilization. UCA1 functions as a molecular sponge for pro-inflammatory miRNAs, such as, miR-143, miR-204, and miR-27b, involved in regulation of cell proliferation, apoptosis, survival and modulate *FOXO1*, *BCL2*, and PI3K/AKT pathway, under cellular stress and viral inflammation.^52–54^ Therefore, the acquisition of novel m⁶A sites may enhance transcript stability and indirectly promote the translation of target genes to alter the inflammatory response. In our study, UCA1 exhibited the highest number of m⁶A sites with high modification probability (>90%) compared to other lncRNAs analyzed, indicating an intense epitranscriptome activity in SARS-CoV-2 infection. The presence of multiple highly methylated residues indicates that UCA1 is a preferential target of m⁶A methylation machinery. This is supported by the demonstration of targeted recruitment of METTL3/14 to specific regulatory RNAs under stress and viral infection, resulting in functional alterations in the stability and translation of these RNAs.^55,56^

### DRACH motif nucleotide bias

DRACH motif analysis revealed subtle alterations in sequence context, which may reflect increased substrate specificity of the methyltransferase complex under infection conditions (**Figure 3**, **Table 3**). Although the GAC core is conserved, an enrichment of A/U nucleotides was observed in the flanking regions in infected cells, particularly among lncRNAs with a higher number of variant m⁶A sites. This shift in sequence profile may reflect a reprogramming of methyltransferase substrate specificity, favoring m⁶A methylation in more permissive sequence contexts. Such changes may be linked to alterations in the activity of cofactors such as WTAP,^4^ and modulation of DRACH preferences by SARS-CoV-2 itself that suggests a viral adaptation mechanism aimed at exploiting the host’s epitranscriptome machinery.^6^

### Intra- and inter-molecular duplexes of lncRNAs and non-canonical base pairs

The present analysis shows the enrichment of m⁶A sites in intra-molecular duplexes, such as lncRNA secondary structure hairpins, and one in an inter-molecular RNA duplex (UCA1-IFNAR2) (**Figure 4** and **Figure 5**). These modifications coincide with observed changes in expression levels of known lncRNA targets, suggesting that the m⁶A modifications interfere with regulatory activity of lncRNAs.^57^ Notably, among the lncRNAs analyzed, only UCA1-001 (ENST00000397381) presented two methylated m⁶A sites (positions 848 and 864) within the predicted region of interaction with its target (in UCA1-001 at 808–870 and IFNAR2-001 at 2853-2900) (**Table 3**, **Figure 4**). This is an exception to the general pattern we observed: lncRNA–target interaction sites tend to avoid DRACH motifs and m⁶A methylation, suggesting a selective pressure against direct methylation in these regions (**Table 3**). Intriguingly, in the interaction regions when a DRACH motif is present in the lncRNA, the corresponding target lacks one, and vice versa, except for UCA1 interaction (positions 2013-2055) with IFNAR1 (positions 59-102) (**Table 3**). Rather than acting directly at the interaction interface, m⁶A modifications appear to exert their effects by modulating the secondary or tertiary structures of lncRNAs.^58^ These structural changes could influence target recognition by altering accessibility and binding. Changes in lncRNA structure associated with altered m⁶A sites can also impact the accessibility to reader proteins that mediate RNA stability. Therefore, no single, overarching rule governs the presence of m⁶A within RNA interaction regions, which suggests a regulatory trend mediated by context-dependent structural modulation, in keeping with the dynamic nature of lncRNA function.

One potential molecular mechanism underlying this phenomenon involves the destabilization of lncRNA duplexes via m⁶A-dependent interference with base pairing. It is well-established that m⁶A disrupts canonical Watson-Crick A-U base pairing by introducing steric hindrance at the *N*⁶ position of adenosine, thereby lowering the thermodynamic stability of RNA duplexes.^59^ Beyond this destabilization, recent structural studies suggest that m⁶A may promote the transient formation of Hoogsteen-like base pairs.^57,60^ Although Hoogsteen conformations are generally disfavored in A-form RNA helices, their transitory occurrence may contribute to local instability within RNA-RNA interaction interfaces. In the context of lncRNA-target recognition, m⁶A-induced switching between canonical and non-canonical base-pairing geometries could reduce the effective residence time of lncRNAs on their mRNA or miRNA targets, weakening or abolishing regulatory interactions. We propose that m⁶A methylation destabilizes canonical A–U base-pairing in hairpins and RNA–RNA duplexes, enabling transient Hoogsteen-like base-pairing. This conformational switch could reduce the stability of lncRNA-target hybrids, altering regulatory efficiency. Such structural shifts may explain the concordant changes in lncRNA methylation and gene expression we observed in SARS-CoV-2-infected cells (**Figure 7**). The non-canonical base pairings render the RNA-RNA hybrids more unstable especially in positions involved in hairpin structures (**Figure 5**). m⁶A disrupts Watson-Crick A-U pairing and can transiently switch to Hoogsteen A-U pairing in A-rich loops^57^ in RNA-RNA hybrids^25^ (**Figure 7**). The m⁶A–Hoogsteen model offers a unifying hypothesis linking lncRNA methylation shifts to altered gene expression, opening new avenues to explore how RNA modifications dynamically modulate RNA–RNA interactomes during viral infection.

**Figure 7.**
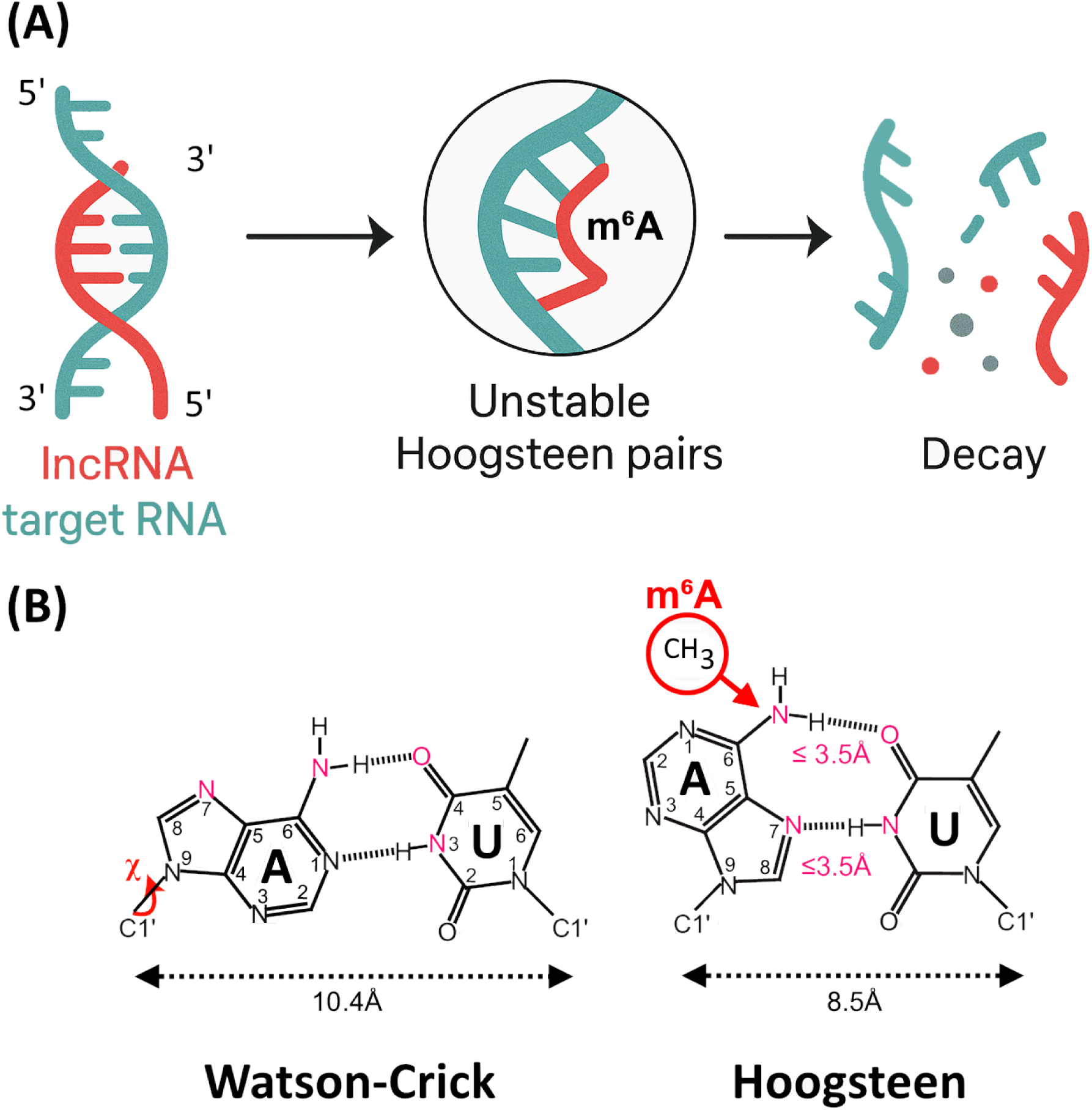
RNA positions with m⁶A and Hoogsteen base pairing. In (A) the target sequence, and surrounding bases, in the target RNA or the target recognition sequence in the lncRNA can be modified by m⁶A because they contain DRACH motifs. In (B) m⁶A methylation can increase the probability of Hoogsteen pairs that turns the hybridization unstable leading to degradation. The addition of a methyl group at the N^6^ position causes the torsion (χ) of the adenine (A) and reduces the width of the RNA-RNA hybrid from 10.4Å to 8.5Å causing its instability. (B) Modified from Zhou et al. (2015)^60^.

Interestingly, among most lncRNA-target interaction regions analyzed, m⁶A-modified DRACH motifs were present but did not participate in base pairing with the complementary strand (**Figure 4**, **Figure 5**). This suggests that these motifs, and the associated m⁶A sites, may be functionally tolerated in other duplex regions but are generally avoided within lncRNA interaction sites. While not conclusive, this observation is consistent with the idea that lncRNA–target interactions may disfavor m⁶A-modified regions, or alternatively, that m⁶A sites are selected against in inter-molecular duplex-forming domains. Notably, DRACH motifs and m⁶A modifications were abundant elsewhere in the lncRNA molecule, suggesting a potential spatial segregation between interaction domains and methylation sites.

### Conclusion

Mapping m⁶A methylation patterns using dRNA-seq data enables single-base resolution of RNA modifications in native molecules without the need for conversion, amplification or antibodies.^5,61^ Remodeling of the host epitranscriptome is documented in mRNAs during SARS-CoV-2 infection^62^ and later validated in immunological models.^63^ By integrating expression profiling, single-nucleotide resolution m⁶A mapping, and DRACH motif analysis in a human pulmonary cell model, we suggest that lncRNAs such as GAS5, NORAD and UCA1 are transcriptionally responsive and preferential targets of the m⁶A machinery. We propose a theoretical model in which SARS-CoV-2 induces m⁶A remodeling of lncRNA. Our analysis suggests that UCA1, particularly, plays a relevant role in SARS-CoV-2 infection response via m⁶A methylation. The m⁶A remodeling in lncRNAs may impact their ability to bind and regulate target RNAs. Transient, unstable Hoogsteen-like base pair geometries can be formed at critical RNA duplex regions that contain m⁶A which possibly affect the stability of RNA-RNA interactions and modify gene expression patterns associated with IFN response. To our knowledge, this is the first comprehensive study to characterize m⁶A dynamics of lncRNAs in SARS-CoV-2 infection in a physiologically relevant *in vitro* model. These findings extend the understanding of host–virus interactions at the epitranscriptome level and highlights the potential of m⁶A-modified lncRNAs as candidates for functional studies and therapeutic targeting.

### Limitations of the study

First, the identification of m⁶A sites and RNA–RNA interactions relied exclusively on *in silico* predictions using dRNA-seq and machine learning models. Second, the analysis was based on Calu-3 cells, a well-characterized but *in vitro* model of pulmonary epithelium. Therefore, the generalizability of these findings to other cell types, tissues, or *in vivo* contexts remains to be clarified. Third, although sequence logo and motif analyses revealed statistically significant remodeling of the DRACH motif landscape, these observations require experimental validation to confirm shifts in substrate specificity of the m⁶A methyltransferase complex during infection. Finaly, the rDNA-seq data was obtained from three independent infections, which were treated as technical replicates in the original study. While the cDNA-seq data were sequenced separately, the RNA samples were pooled for dRNA-seq due to multiplexing limitations in dRNA-seq kits (Oxford Nanopore Technologies). As a result, while dRNA-seq reflects the average behavior across infections, this pooling precludes replicate-aware statistical analyses, such as differential methylation testing with variance estimates. The focus of this study is to infer average changes in lncRNA expression and m⁶A patterns, and in that context, the pooled data remain valid for global trend analysis and hypothesis generation.

## RESOURCE AVAILABILITY

### Lead contact

Requests for data, further information and resources should be directed to and will be fulfilled by the lead contact, Marcelo R. S. Briones.

### Materials availability

This study did not generate new unique reagents.

### Data and code availability

The methylation workflow produced in this work is available on https://github.com/CaioCCTI/lnc-team-m6A/blob/main/

## ACKNOWLEDGMENTS

We thank Marcello Ramon for technical assistance. This work was supported by grants from Fundação de Amparo à Pesquisa do Estado de São Paulo (FAPESP, Brazil), grant 20/08943-5 to M.R.S.B. and F.A. and Conselho Nacional de Desenvolvimento Científico e Tecnológico (CNPq, Brazil), grant 311154/2021-2 to M.R.S.B.

## AUTHOR CONTRIBUTIONS

Conceptualization, M.R.S.B., C.M.P. and N.M.; methodology, C.M.P., C.O.C., F.A. and M.R.S.B; performed the experiments, C.M.P. and C.O.C.; analyzed the data, C.M.P., C.O.C., F.A. and M.R.S.B.; writing—original draft preparation, C.M.P., C.O.C. and M.R.S.B.; writing—review and editing, C.M.P., C.O.C., F.A., N.M. and M.R.S.B. All authors have read and agreed to the published version of the manuscript.

## DECLARATION OF INTERESTS

The authors declare no competing interests.

## METHODS

Detailed methods are provided in the online version of this paper and include the following:

## KEY RESOURCES TABLE

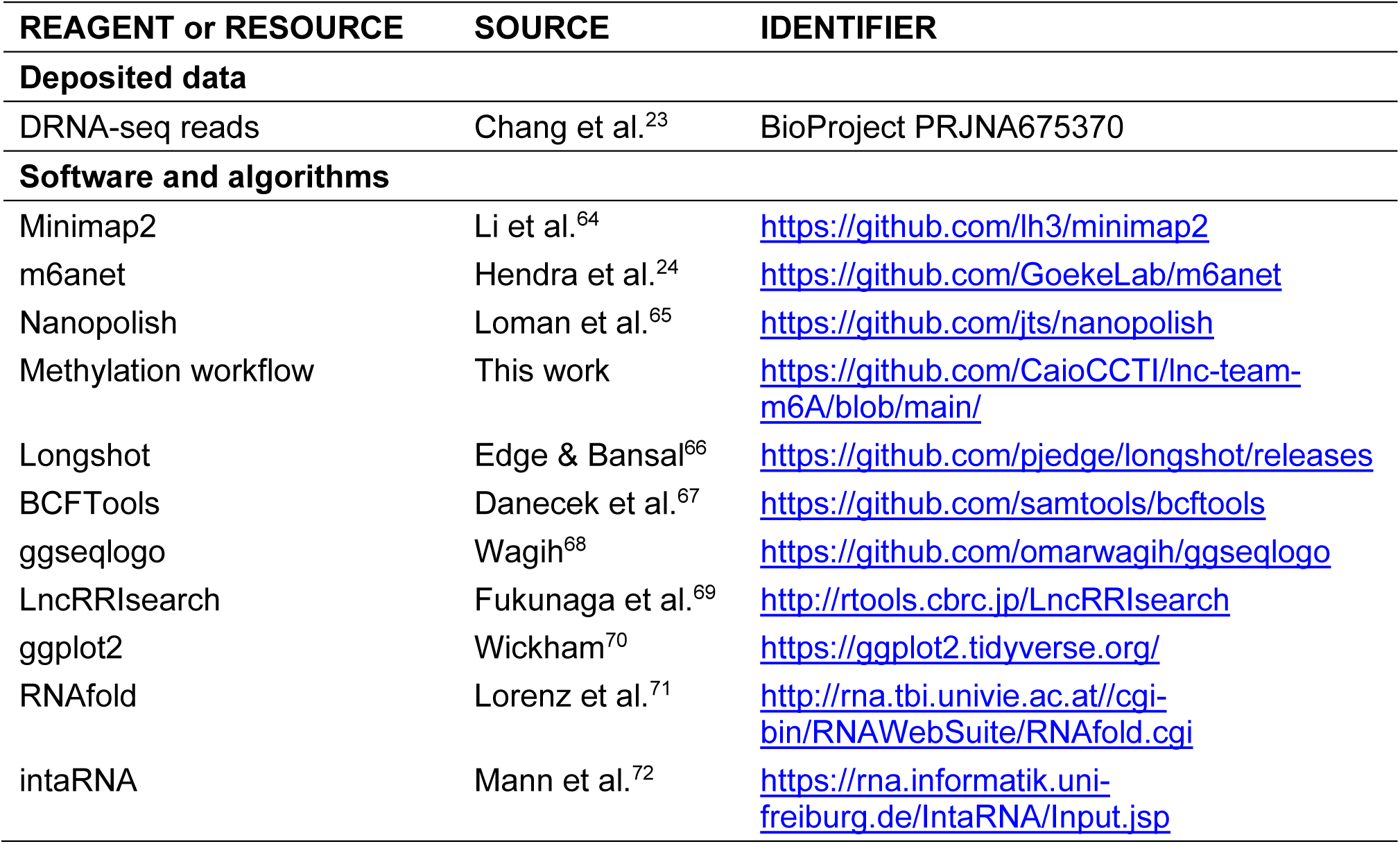

## EXPERIMENTAL MODEL AND SUBJECT DETAILS

RNA sequences (dRNA-seq reads) from SARS-CoV-2 infected and uninfected Calu-3 cells were obtained from Chang et al. (2021)^23^. Calu-3 cells were infected with SARS-CoV-2/Australia/VIC01/2020, NCBI: MT007544.1. These cells are widely used as an *in vitro* model for studying respiratory viral infections, particularly coronaviruses (SARS-CoV, SARS-CoV-2, MERS-CoV) and influenza viruses, due to their high expression of angiotensin-converting enzyme 2 (ACE2), the primary receptor for SARS-CoV-2 entry^73,74^. Calu-3 cells were infected with SARS-CoV-2 for 48 hours, followed by total RNA extraction and dRNA-seq. Reads from control and infected Calu-3 cells were obtained from SRA BioProject PRJNA675370, SRA Accession #SRR13089335 of SARS-CoV-2 infected Calu-3 RNA reads and #SRR13089334 uninfected Calu-3 RNA reads.^23^ The data is from three separate infections of the same cell line performed at the same time and are, therefore, technical replicates rather than biological replicates. While replicates were sequenced separately for cDNA-seq, they were pooled prior to dRNA-seq due to technical limitations in multiplexing with Oxford Nanopore Technologies (ONT) sequencing kits.

## METHOD DETAILS

### Contig assembly by mapping to references

For the identification of lncRNAs of interest in response to SARS-CoV-2, we initially conducted a search in the NCBI (National Center for Biotechnology Information) database using specific terms related to viral infection and lncRNA expression. After selecting the relevant, we downloaded the records in FASTA format, ensuring the preservation of structure and integrity. After downloading it in FASTA format, we used Minimap2 to map the dRNA-seq reads to the lncRNA reference sequences^64,75^. This mapping allowed us to identify the presence and potential variations in lncRNA expression in response to viral infection, providing insights into their possible role in regulating the cellular response to the virus. The Minimap2 mapping generated BAM files for posterior analyses. The expression levels were estimated as fold change (FC) from the number of reads mapped to each lncRNA in infected and uninfected cells using the formula:

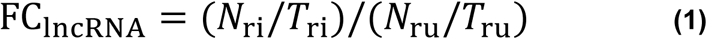

where *N*_ri_= number of mapped reads of each lncRNA in infected cells, *T*_ri_=total mapped reads in infected cells (1,055,956), *N*_ru_= number of mapped reads of each lncRNA in uninfected cells and *T*_ru_= total reads in infected cells (940,040).

### Methylation analysis

From the BAM files and the fast5 files from the Nanopore sequencing the “index” and “eventalign” modules from Nanopolish (v0.14.1.) were utilized for the signal segmentation step, commonly referred to as “squiggling.” The segmented raw signals obtained in this step were stored in the eventalign output file and subsequently pre-processed using the m⁶A net-dataprep module. The detection of m⁶A modifications within DRACH motifs was performed via the m⁶A net-run_inference algorithm, both of which are implemented in m6anet (v2.1)^24^. The complete workflow, including the custom scripts used for data processing and analysis, is available in the research group’s GitHub repository: https://github.com/CaioCCTI/lnc-team-m6A/blob/main/README.md#introduction.

### Analysis of DRACH motifs

For detailed inspection of methylated DRACH motifs - checking for information bias – the consensus sequences were obtained from the variation calling step. This procedure was performed with Longshot (v.0.4.1), with default settings for long reads, and establishing a minimum threshold of read depth for 30x coverage to accept the SNV occurrence^66^. The resulting VCF files obtained in this step were used to generate consensus by the “bcftools consensus” - BCFTools (v.1.15)^76^. All methylated DRACH motifs were subsequently extracted and flanked by five nucleotides on each side to enable sequence alignment and stacking. This procedure was implemented using a custom computational pipeline developed in Python (version 3.13.2)^77^.

To minimize the occurrence of m⁶A sites that may represent false positives in Calu-3 cell samples (presence of methylated m⁶A sites with a reduced probability of predictions as a function of many transcripts/reads) and that could add noise to the analysis, the inspection of the DRACH motifs was performed using m⁶A sites with ≥ 0.6 probability of modification threshold as calculated by m6anet^24^. To reduce the occurrence of false negatives in SARS-CoV-2 with fewer reads, and a smaller genome size, the methylation probability threshold was set at ≥ 0.6. Because some DRACH motifs (including 5 nucleotides at each end) can be located at the coordinate ends of cellular transcripts and viral RNAs, resulting in truncated sequences, the removal of these sequences was necessary. The analysis of nucleotide biases within DRACH motifs was conducted using the ggseqlogo package (v.0.2) implemented in R (version 4.4.2)^68^.

### • Statistical tests of differential methylation

To compare the distribution of methylated sites per transcript in sequencing reads from control and infected cells, we performed the Wilcoxon-Mann-Whitney (WMW) test, as implemented in R version 4.4.2 within the base R package^78,79^. This nonparametric test is widely used to assess differences between two independent groups when their distributions do not meet the assumptions of normality. Under the null hypothesis (H₀), the distributions of both groups are assumed to be identical, whereas the alternative hypothesis (H₁) posits that the distributions differ significantly, specifically by detecting a difference in their medians.

To apply the WMW test to our data, we first gathered all predictions generated by m⁶A net and stored them in tables, which were subsequently imported into R as data frames and labeled as “uninfected” and “infected.” We then applied a probability threshold (modification probability ≥ 0.6) to filter out sites with low confidence in m⁶A modification. The primary variable chosen to compare the methylation distribution between the two groups was the number of reads per site (*n* reads). Since the number of reads per site is not standardized by default, data normalization was required. For this, the number of reads per site in each group was divided by the total number of reads in that group using the equation below. After normalization, the data underwent a negative logarithm transformation (−log) to enhance visualization consistency across samples:

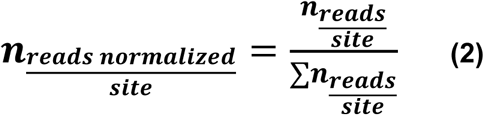

To visualize the distributional differences between groups, we generated violin plots using the ggplot2 package (version 3.5.1) in R^70^. Violin plots provide comprehensive graphical representation of data distribution by combining a box plot with a kernel density estimate, enabling the visualization of both summary statistics and the underlying probability density function.

### Prediction of lncRNA–mRNA interactions and integration with m⁶A methylation analysis

Interactions between long non-coding RNAs (lncRNAs) and coding mRNAs were predicted using the LncRRIsearch platform (http://rtools.cbrc.jp/LncRRIsearch), which allows for the systematic identification of RNA–RNA interactions based on sequence complementarity and structural accessibility^69^. This methodology considers thermodynamic parameters and RNA secondary structure to predict potential base-pairing regions between transcripts. The lncRNAs GAS5, NORAD, and UCA1, in their reference isoforms, were used as input for the interaction predictions. Target mRNAs were selected based on their relevance to antiviral response, IFN signaling, and inflammatory regulation based on experimental evidence related to SARS-CoV-2 infection.^80^

For each predicted interaction, nucleotide positions corresponding to the interaction region between the lncRNA and mRNA were extracted. These data were then integrated with epitranscriptomic profiles generated through dRNA-seq (Oxford Nanopore Technology) and analysis with m6anet for the identification of m⁶A sites. The relationship between methylation and RNA–RNA interactions, we looked at overlaps between predicted binding regions and canonical DRACH motifs, which define the consensus sequence recognized by the m⁶A methyltransferase complex. Only interactions coinciding with high confidence m⁶A sites (as predicted by m6anet) were retained for downstream analysis. Methylation events were categorized according to their genomic context: m⁶A in the lncRNA, when the modification occurred within the lncRNA region involved in the interaction, m⁶A in the target gene, when located within the interacting region of the target mRNA and m⁶A in both, when m⁶A sites were detected in both interacting partners. The lncRNA-SARS-CoV-2 interactions were predicted using intaRNA.

## SUPPLEMENTAL INFORMATION

Supplemental information can be found online at http.

## Notes

### Competing Interest Statement

The authors have declared no competing interest.

